# SNPstar: A Web Server Linking Allelic Variation to Protein Structure and Function in *Arabidopsis thaliana*

**DOI:** 10.64898/2026.07.16.738854

**Authors:** Benjamin Schmidt, Laura Pilgram, Steve Babben, Jana Trenner, Selma Gago-Zachert, Francesco Pezzini, Christian Tüting, Sven-Erik Behrens, Muhammad Tahir, Jan Grau, Georg Künze, Panagiotis L. Kastritis, Ivo Grosse, Marcel Quint

## Abstract

Understanding how natural genetic variants affect protein structure and function is central to plant biology. The 1001 Genomes Project has catalogued millions of single nucleotide polymorphisms (SNPs) across more than a thousand *Arabidopsis thaliana* accessions, offering an unprecedented opportunity to relate sequence variation to three-dimensional protein structure and population context. Yet, realizing it requires tools that integrate these scales in one accessible framework. Here we present SNPstar, a web server that links allelic variation in *A. thaliana* to AlphaFold3-predicted structures through a gene-centric, interactive interface. SNPstar annotates each variant with descriptive features, thermodynamic stability estimates, protein domain context, and genome-wide association results, and computes haplotypes and proteotypes that group accessions by shared DNA or protein sequence. Researchers can characterize variants, visualize their structural context, map their geographic distribution, and prioritize accessions for experimental validation without local computational infrastructure. We demonstrate SNPstar with two case studies. The first recapitulates known loss-of-function variation in the cadmium transporter *HMA3*, validating that SNPstar prioritizes functionally consequential alleles. The second uses SNPstar-defined proteotypes to identify an N-terminal SNP combination in *ARGONAUTE 2* that distinguishes accessions differing in *in vitro* siRNA-directed target cleavage, linking protein-coding variation to a measurable molecular phenotype. SNPstar thus helps translate natural variation into mechanistic insight.

## Introduction

The integration of genomic variation with protein structure and function has emerged as a critical frontier, particularly in plant biology, which still lags behind in terms of experimentally generated protein structures (Burley et al. 2025). High-throughput sequencing initiatives in plants, beginning with *Arabidopsis thaliana* (Cao et al. 2011) and the landmark 1001 Genomes Project (The 1001 Genomes Consortium 2016), have generated unprecedented catalogs of genetic variation. Although these data sets have identified millions of variants, their analysis presents significant challenges, particularly when identifying variants that impact protein structure and function. Understanding how sequence polymorphisms affect protein function provides essential insights into both fundamental biology and crop improvement. *Arabidopsis thaliana* serves as an ideal model organism for these studies because of its compact genome, high proportion of coding sequences, and extensive naturally occurring genetic variation. These features allow researchers to link genotypes to phenotypes at the species-level that can potentially be translated to agronomically important crops.

Between genotype and phenotype, genetic variation manifests at multiple molecular levels: haplotypes at the DNA level (defined by both synonymous and non-synonymous SNPs), proteotypes at the amino acid sequence level (reflecting only non-synonymous SNPs), and proteoforms, which additionally incorporate post-translational modifications of the proteotype. Whereas haplotypes and proteotypes are directly encoded in the genome, proteoforms arise post-translationally and can therefore be addressed only indirectly in genome-based tools such as SNPstar (see use case below).

Recent advances in protein structure prediction - through methods like AlphaFold (Jumper et al. 2021; Abramson et al. 2024), ESMFold (Lin et al. 2023), RoseTTAFold (Baek et al. 2021; Krishna et al. 2024; Corley et al. 2025) and others (Discovery et al. 2024; Wohlwend et al. 2025) - have created opportunities to contextualize genetic variants within three-dimensional protein structures. The convergence of these computational approaches, recognized by the 2024 Nobel Prize in Chemistry, has established the technical foundation for more sophisticated variant interpretation. What remains is to bring this structural perspective together with population-scale variant data in a single analytical environment. Several established resources each cover important parts of this landscape (Table S1). Polymorph 1001 (1001 Genomes Project 2024) and ViVa (Hamm et al. 2019) provide rich access to genetic variant and population data; AraGWAS (Togninalli et al. 2017) connects variants to phenotypes through genome-wide association studies; UniProt (The UniProt Consortium 2025) brings together protein structures, curated variants, and functional annotations; and Ensembl Plants (Yates et al. 2021a) has begun to combine protein structure with variation data. Each excels in its respective domain, but a unified, gene-centric view that links population-level variation to predicted protein structures - and adds proteotype-level analysis on top - has so far been missing. SNPstar is designed to fill this niche, allowing researchers to investigate structure–function relationships of natural variants within a single interactive interface rather than transferring findings manually between platforms.

Here, we present this unified framework in detail. SNPstar enables researchers to simultaneously view variants in their sequence context, evaluate their population distribution across accessions, and visualize their spatial positioning and effects within AlphaFold- and FoldX-predicted protein structures (see Table S1 for feature comparison with published tools). It integrates variant data, protein domain information, GWAS results, and structural models of different proteotypes, allowing researchers to address biological questions about SNPs that span scales from global population structure down to the positioning of specific side chains within a proteotype.

The main SNPstar genetic screening workflow supports gene-centric investigations, enabling researchers to analyze the potential structural and functional effects of individual variants, haplotypes, and proteotypes. By streamlining the assessment of SNP effects on protein structure and function, SNPstar helps researchers prioritize variants and proteotypes for experimental validation.

The paper is structured as follows. We first describe the data underlying SNPstar and provide a brief overview of its implementation, and then outline its key functionalities within a gene-centric workflow. A detailed case study based on published data illustrates this workflow in practice. We then apply SNPstar to a previously uncharacterized molecular phenotype, natural variation in siRNA-directed target cleavage (slicing) by ARGONAUTE 2 (AGO2) across Arabidopsis accessions, and close with a discussion of limitations and future developments.

## Methods

### Implementation and Data

SNPstar’s architecture comprises three components: a variant-processing pipeline, a set of precomputed protein structure models, and a web application stack; described in turn below.

### Data Processing Pipeline

The SNPstar setup pipeline processes genomic variation data from the 1001 Genomes Project for *Arabidopsis thaliana* (The 1001 Genomes Consortium 2016) using the Araport 11 annotation (Cheng et al. 2017). This Snakemake-based workflow (Mölder et al. 2025) systematically annotates variants through SNPEff (Cingolani et al. 2012) to categorize and predict functional impacts. Python scripts then derive haplotypes and proteotypes from SNPEff output. For a given gene, all accessions that share the same set of SNPs within that gene - i.e., the same DNA sequence - have the same haplotype. Proteotypes are the protein-sequence-level equivalent: they group all accessions that share the same amino-acid sequence for a given gene. Protein domain information is integrated from InterProScan (Jones et al. 2014), while 15,839 significant associations (p < 10^-4^ according to a permutation analysis) are incorporated from the AraGWAS catalog (Togninalli et al. 2017). AraGWAS standardizes GWAS results across the *A. thaliana* research community. Additional GWAS analyses for climate factors were performed as described by Ferrero-Serrano and Assmann (2019), using the software GEMMA (Zhou et al. 2013). Each climate factor was treated as a phenotype in a separate GEMMA run. Accessions with missing phenotype values were excluded prior to analysis. Minor allele frequency (MAF) and linkage disequilibrium (LD) thresholds in GEMMA were set to 1% and 0.8, respectively. The resulting *p*-values were corrected for multiple comparisons using the Benjamini–Yekutieli method.

### Protein Structure Models

SNPstar provides protein structural models for the standard *Arabidopsis thaliana* proteome and for proteotypes with an allele frequency above 1%. Structural models are available in SNPstar for all annotated protein sequences in all accessions, including those based on alternatively spliced transcripts. Structure predictions were made using AlphaFold3 (v3.0.0) (Abramson et al. 2024), with multiple sequence alignments (MSAs) generated via the MMseqs2-based ColabFold v1.5.5 workflow (Mirdita et al. 2022), searching the UniRef30 (2023_02) and ColabFold environmental (2021_08) databases, and using AlphaFold’s template mode with default settings. Of the five models generated per sequence, the top-ranked model based on AlphaFold’s ranking score metric is used for visualization in the web interface. The proteotype models were generated by applying FoldX 5.0 (Delgado et al. 2025) to the AlphaFold3 Col-0 models. FoldX’s computational efficiency permits the modeling of several hundred thousand proteotypes while providing per-model stability predictions. Each AlphaFold3 model was first refined with the RepairPDB module in FoldX, followed by modeling of the particular amino acid mutations of each proteotype using the BuildModel module with five iterations. For proteotypes containing insertions or deletions relative to the Col-0 reference, the reference sequence was first shortened or extended accordingly and remodeled with AlphaFold3 before introducing the point mutations with FoldX. The proteotype model with the lowest total energy was selected for visualization. In addition, FoldX and ThermoMPNN (Delgado et al. 2025, Dieckhaus et al. 2024) energetic stability predictions (ΔΔG) are available for all proteotype models and can be visualized in the context of the protein structure.

### Application Stack

For the webserver, the structural data, variant data, annotations, and computed features described above are imported into our database. As a database management system, SNPstar uses MongoDB (MongoDB Inc. 2024) with optimized query indices for efficient retrieval. A Python Flask application handles query processing, dynamically computing results for user-selected accession sets and genes. This implementation enables rapid filtering and subset analysis across large variant datasets. The web server employs Gunicorn (Gunicorn Team 2024) to handle concurrent connections reliably, ensuring consistent performance under varying user loads.

The user interface integrates specialized JavaScript libraries to provide a responsive analytical environment: nextProt (Zahn-Zabal et al. 2019) components for sequence feature visualization, Webix (XB Software Ltd. 2024) for interactive data tables, Leaflet with OpenStreetMap (OpenStreetMap Contributors 2024) for geographic distribution mapping, and Mol* (Sehnal et al. 2021) for three-dimensional protein structure rendering. This integrated visualization approach enables researchers to explore connections between sequence variation, protein structure, and geographical distribution through multiple linked views.

### Availability

SNPstar is freely available at http://snpstar.informatik.uni-halle.de. Source code is maintained at https://github.com/GrosseLab/SNPstar, and complete datasets are accessible at https://snpstar.informatik.uni-halle.de/arabidopsis_thaliana#about. In addition, the workflows, input and output data are provided as an Annotated Research Context (ARC) (Weil et al. 2023) following the FAIR principles (Wilkinson et al. 2016) at https://git.nfdi4plants.org/snp2prot/public/snpstar. Using the available codebase and data integration pipeline, SNPstar can also be deployed locally - for example, to analyze proprietary variants, experimental phenotypes, or custom structural models within secure computing environments.

### Experimental validation of AGO2 proteoform activity

#### Plasmid construction

Inserts containing the complete *AGO2* coding sequence were generated by RT-PCR. cDNA was synthesized with an oligo(dT)18 primer (Thermo Fisher Scientific, Dreieich, Germany) and RevertAid reverse transcriptase (Thermo Fisher Scientific) from total RNA extracted from seedlings of the Col-0 (ABRC ID CS76778) and the BRR4 (ABRC ID CS78943) accessions. Amplicons were generated with primers RR952 (5′-caccATGGAGAGAGGTGGTTATCGAGG-3′) and RR953 (5′-TCAGACGAAGAACATAACATTCTCAAG-3′) and Phusion High-Fidelity DNA polymerase (Thermo Fisher Scientific), cloned into pBluescriptKS(+) (Stratagene, Agilent Technologies, Santa Clara, CA, USA) linearized with *Eco*RV, and subcloned into the pSPLF2 plasmid (Schuck et al. 2013) - a modified variant of the pSP64-Poly(A) vector (Promega, Madison, WI, USA) - digested with *Sma*I and *Xba*I. The FLAG-tag sequence was added by ligating annealed oligonucleotides encoding the tag at the 5′ end of the insert-containing pSPLF2 vector.

#### Cell culture and preparation of BYL

*Nicotiana tabacum* BY-2 cells were cultured at 23 °C under continuous agitation in Murashige and Skoog (MS) medium (Duchefa Biochemicals, Haarlem, Netherlands). Cytoplasmic extract (BYL) was prepared from evacuolated cells as described (Komoda et al. 2004; Gursinsky et al. 2009).

#### In vitro synthesis of RNA transcripts

RNA was synthesized by *in vitro* transcription following standard procedures. FLAG-tagged *AGO2* mRNAs were synthesized from *Smi*I-digested plasmids using SP6 RNA polymerase (Thermo Fisher Scientific) and the monomethylated cap analogue m7GpppG (Jena Bioscience, Jena, Germany). Antisense firefly luciferase RNA was generated from the pSP-luc(+) plasmid (Promega) digested with *Hin*dIII using the TranscriptAid T7 High Yield transcription kit (Thermo Fisher Scientific). Fluorescently labelled GFP target RNA was generated from 500 ng of a purified PCR fragment following standard procedures in the presence of 1 mM ATP and CTP, 250 µM GTP, 150 µM UTP, 0.8 mM m7GpppG cap analogue, 50 µM UTP-Atto 680 (Jena Bioscience) and 60 U of T7 RNA polymerase (Thermo Fisher Scientific). Transcripts were purified with the Monarch Spin RNA Cleanup Kit (New England Biolabs, Frankfurt am Main, Germany).

#### In vitro slicer assay and catalytic activity determination

AGO2/RISC loaded with a specific siRNA was generated as described (Knoblich et al. 2025). After 3 h incubation at 25 °C, a 10 µL aliquot was taken for protein analysis, and 10 µg of antisense firefly luciferase mRNA (competitor for nonspecific RNases) and 312.5 fmol of the fluorescently labelled GFP target RNA were added. Aliquots were taken at defined time points and the reaction stopped with 1% SDS. RNA was isolated, purified, precipitated and resolved on 15% denaturing (8 M urea) polyacrylamide gels. Gels were scanned on an Odyssey CLx imager (LI-COR Biosciences, Lincoln, NE, USA) and band intensities quantified with Image Studio v5.2 (LI-COR). Degradation rate constants were determined from these values in KaleidaGraph (v4.5.2 for Windows, Synergy Software, Reading, PA, USA). Values were obtained from at least three independent assays.

## Results and Discussion

### Usage

The genetic screening workflow (Figure 1) is designed to help researchers identify non-synonymous SNPs and proteotypes most likely to affect protein structure and function, and select promising candidates for experimental validation.

**Figure 1.**
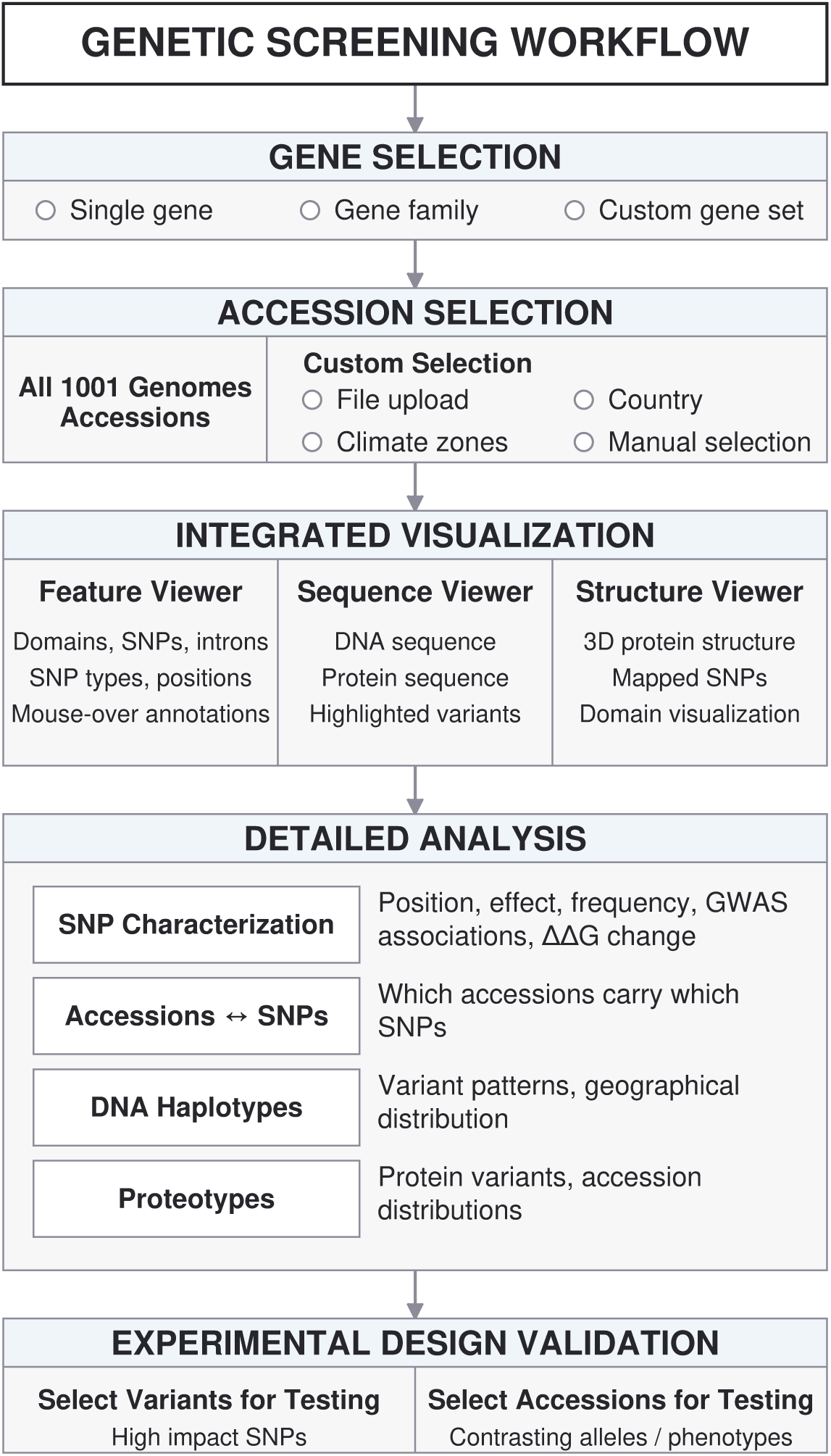
Overview of SNPstar’s genetic screening workflow. The user first selects one or several target transcripts (single gene, gene family, or custom set) and a set of accessions (all 1001 Genomes accessions or a custom subset defined by file, country, climate zone, or manual selection). Integrated visualization then provides parallel views of the feature viewer (domains, SNPs, introns), sequence viewer (DNA and protein sequence), and structure viewer (3D protein structure with mapped SNPs). Detailed analysis is supported by four tables - SNP characterization, SNP–accession mapping, DNA haplotypes, and proteotypes - that surface position, effect, allele frequency, GWAS associations, predicted ΔΔG, and accession distribution. Together, these views support the selection of variants for testing and of accessions carrying contrasting alleles for experimental validation.

The web tool includes an automatic ‘*Getting Started*’ tutorial (Figure S1) on the initial ‘Select transcripts and accessions’ page that guides users through the most important SNPstar features. These features are organized in a central tab structure that includes a track-based genome browser, a DNA and amino acid sequence viewer, a protein structure viewer, and four tabular views: a comprehensive SNP characterization table, a SNP-accession table, a haplotype table, and a proteotype table. A more detailed tutorial covering all tabs and broader SNPstar functionality is available through the ‘*Detailed Tutorial’* icon.

### Gene-centric workflow

The gene-centric workflow allows users to retrieve SNP information for one or more target transcripts (a transcript here refers to a specific splice variant of a gene; the primary transcript ATxGxxxxx.1 is the default). The single-transcript mode is the primary use case, while the multi-transcript mode supports systematic exploration across gene families: SNPstar enables bulk retrieval for a user-defined transcript set and helps identify candidates with particularly informative variant profiles or structural alterations. Both modes can be run on the full set of 1001 Genomes accessions or on user-defined subsets; for example, accessions from particular climatic or geographic regions. Users begin by selecting the accessions of interest, followed by the candidate transcript(s) to analyze.

#### SNP overview in sequence context

After the user has selected a target transcript and a set of accessions, SNPstar provides a track-based genome browser overview of the SNPs (Figure 2A) alongside a sequence-based visualization of the SNPs in the target DNA or protein sequence (Figure 2B). The genome browser displays the DNA-level context, including UTRs, introns, and the coding sequence, while the sequence viewer shows both the DNA and the protein sequence. All views also highlight the locations of conserved protein domains identified with InterProScan, and SNPs are colored by category, with non-synonymous SNPs visually emphasized.

**Figure 2.**
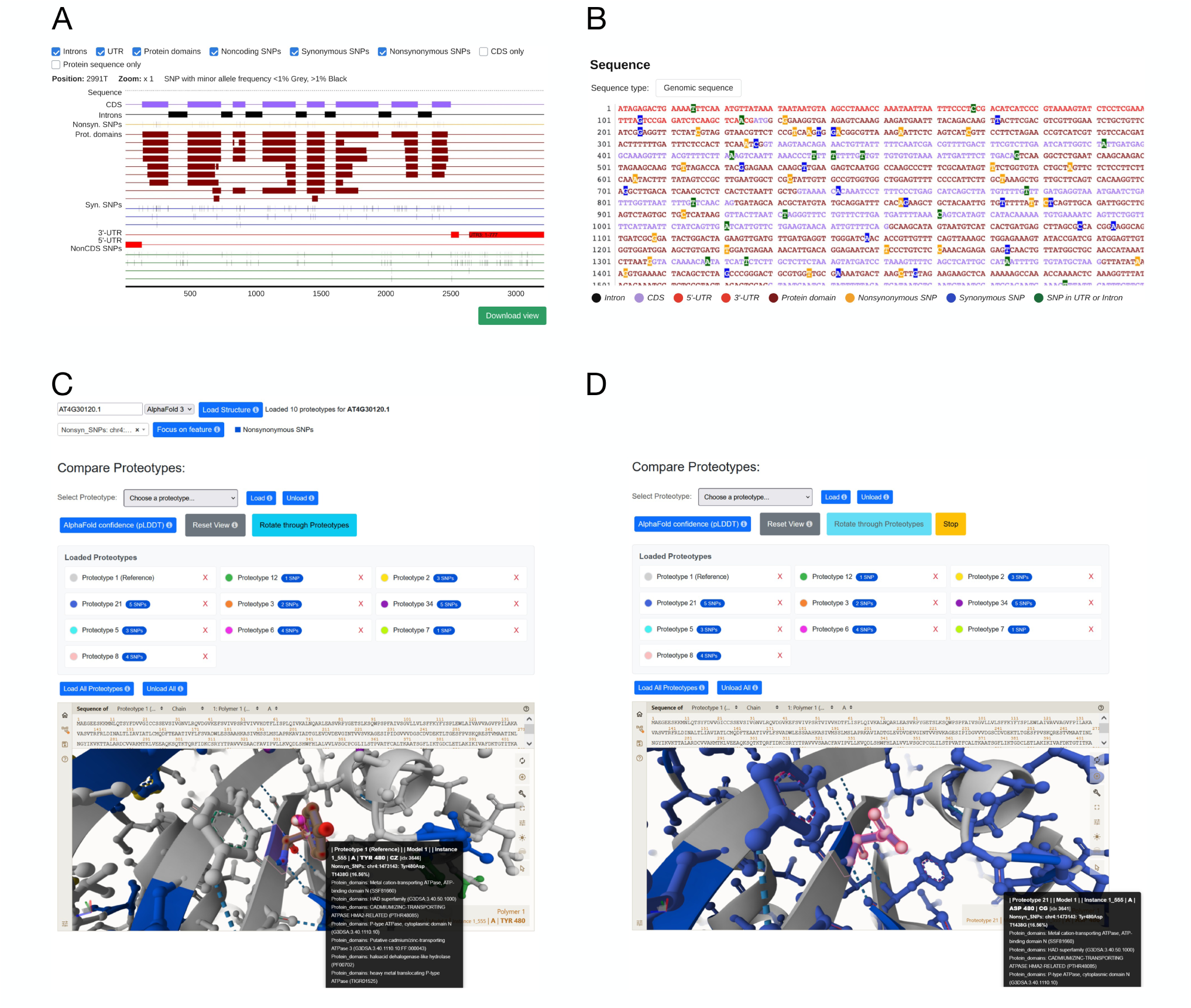
Main SNPstar interface components. (A) Feature viewer showing the genomic context of HMA3 (AT4G30120), with non-synonymous SNPs, synonymous SNPs, non-coding SNPs, UTRs, introns, the CDS, and InterPro protein domains as toggleable tracks. (B) Sequence viewer showing the DNA and protein sequence, with SNPs color-coded by category (non-synonymous, synonymous, UTR/intron) and protein domain regions highlighted. (C) Structure viewer with multiple HMA3 proteotype models loaded simultaneously; non-synonymous SNP positions are highlighted, and mouseover reveals variant details and overlapping protein domains. (D) Structure viewer with a single proteotype (here proteotype 21) brought to the foreground using the rotate-through-proteotypes mode; the variant residue at position 480 is highlighted in pink. Detailed walkthroughs of each tab are available through the “Detailed Tutorial” link in the interface.

Users can (zoom in and) mouse over the non-synonymous SNPs in the genome browser to display the specific amino acid substitution and the allele frequency, and clicking a SNP opens the SNP characterization table (Figure 3A) directly at that variant. For example, a user could identify a non-synonymous SNP in a protein domain, observe a relatively high allele frequency through the mouseover (a possible signature of selective pressure) and then proceed directly to the structure viewer (Figure 2C,D) for closer inspection.

**Figure 3.**
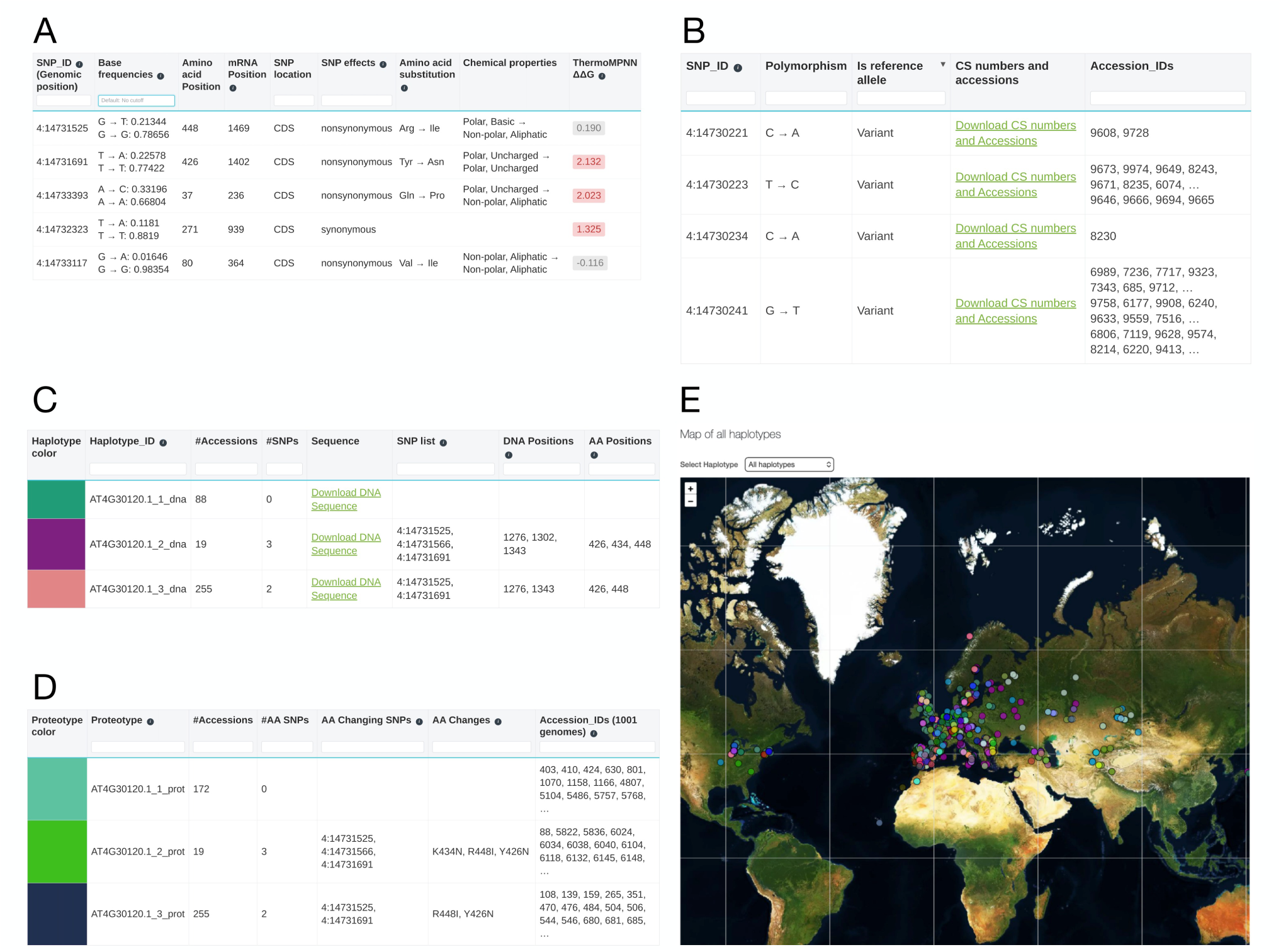
Tabular analysis views in SNPstar. (A) The SNP characterization table provides per-variant annotations including SNP identifier (genomic position), base frequencies, mRNA and amino-acid position, SNP location (CDS/UTR), functional effect, SNPEff impact category, amino-acid substitution with associated change in chemical property class, predicted ΔΔG from ThermoMPNN and FoldX, impact annotation (not shown), InterPro domain assignment (not shown), AraGWAS top hits (not shown, Togninalli et al. 2017), and climate-factor GWAS results (not shown). (B) The SNP–accession table maps each variant to the accessions carrying it, listed by CS number, common name, and 1001 Genomes ID. (C) The haplotype table groups accessions by shared DNA sequence and lists the haplotype ID, frequency, sequence download link, SNP set, and underlying DNA and amino-acid positions for each. (D) The proteotype table groups accessions by shared protein sequence and lists the proteotype ID, frequency, number of amino-acid-changing SNPs and their effects, and the accessions belonging to each proteotype. (E) Depending on the number of accessions selected in the initial ‘Select transcripts and accessions’ tab, geographical origins of haplo- or proteotypes are displayed on a zoomable world map.

#### SNP characterization and functional annotation

In the SNP characterization table (Figure 3A), each variant is annotated along four dimensions: (i) descriptive features - SNP identifier (genomic position), base frequencies, mRNA position, amino-acid position, and SNP location (CDS, 5′ UTR, or 3′ UTR); (ii) functional consequence - SNP effect (synonymous, non-synonymous, or non-coding), SNPEff impact annotation (HIGH, MODERATE, LOW, or MODIFIER), amino-acid substitution, and the resulting change in chemical property class; (iii) predicted structural impact - per-variant ΔΔG values from ThermoMPNN and FoldX (the predicted change in protein stability caused by a mutation; ΔΔG > 0: the mutation is destabilizing; ΔΔG < 0: the mutation is stabilizing); and (iv) functional and population-level context - InterPro domain assignment (ID and description), top hits from the AraGWAS catalog, and climate-factor GWAS results computed as described in the Methods.

Users can filter and prioritize variants based on multiple criteria (for example, allele frequency, genomic localization, predicted functional impact, or domain context) to identify candidates worth further inspection. The interface is also linked bidirectionally to the other analysis tabs: clicking a SNP identifier opens the SNP–accession table (Figure 3B) at the corresponding variant, and the haplotype and proteotype tables (Figure 3C, D) and structure viewer (Figure 2D) are reachable in a single step from the same row.

#### SNP visualization in structural context

The structure viewer (Figure 2D) is the structural counterpart to the sequence- and table-based views and lets users inspect the predicted three-dimensional structures of the Col-0 reference and of all proteotypes with an allele frequency above 1%. The underlying repository contains more than 330,000 structures: AlphaFold3 models for the Col-0 reference proteome and FoldX-derived models for each frequent proteotype (see Methods). Together, these models support detailed analysis of how proteotypes differ from one another and what effect individual SNPs, or combinations of SNPs in the same proteotype, exert on the local structural context.

By default, the viewer shows the Col-0 reference structure with all non-synonymous SNP positions and InterPro protein domains highlighted, helping users assess where variants sit relative to functional regions. Mouseover at a SNP position displays the amino-acid substitution, allele frequency, and any overlapping domain annotations, while the structure can be rotated and zoomed using standard Mol* controls (Sehnal et al. 2021). For instance, users can examine the surface accessibility of a variant site and its spatial proximity to neighboring residues, including residues at known protein–protein interaction interfaces.

A search field lets users locate specific SNPs or domains and zoom directly to their position in the structure. The viewer also displays AlphaFold’s predicted local distance difference test (pLDDT) metric as a color-coded per-residue confidence score, so users can verify whether the structure at the position of interest is reliable before drawing biological conclusions.

To compare proteotypes, users can load any subset of the available proteotype models from a selection field that also lists the SNPs each proteotype carries. Each loaded proteotype is assigned a color, and the variant residues are rendered in ball-and-stick representation on top of the cartoon backbone. A “rotate through proteotypes” mode cycles which proteotype is shown in the foreground, making structural differences between them easier to perceive (see Figure 6 for a worked example in the case study).

#### Accession, haplotype, and proteotype Identification

To facilitate experimental design and validation, SNPstar enables efficient identification of accessions carrying specific variants of interest. The SNP characterization table (Figure 3A) is connected to a SNP-accession table (Figure 3B) that maps each SNP to the accessions carrying it, listed by CS number, common name, and 1001 Genomes ID. This allows researchers to identify variant-carrying accessions already in their collection or to select accessions to order from stock centers via the CS number.

Two further tables provide the haplotypes (DNA-level groupings; Figure 3C) and proteotypes (amino-acid-level groupings; Figure 3D) for the selected gene. Each row lists the accessions belonging to that haplotype or proteotype, its frequency within the population, the underlying SNP set, and any associated start- or stop-codon variation. A linked Leaflet-based world map (Methods) further shows the geographic distribution of each haplotype and proteotype, supporting the selection of accessions from defined climatic or geographic ranges (Figure 3E).

Together, the SNP characterization, structural, and accession-level views give users the information they need to design targeted follow-up experiments. This could be, for example, phenotypic comparisons between accession groups carrying contrasting haplotypes, or site-directed mutagenesis followed by assays of protein stability, post-translational modifications, or protein-protein interactions. Before demonstrating a working example, we first step back to the genome-wide view of the integrated dataset.

### Global properties of variants and structural models

To explore the global properties of non-synonymous SNPs on a genome-wide scale, beyond the gene-centric analyses that are SNPstar’s main purpose, we combined the 1001 Genomes variant catalog with AlphaFold3 structural models and FoldX/ThermoMPNN stability estimates. This integration revealed two general properties of the dataset that frame the case studies below.

We first examined how non-synonymous SNPs are distributed across regions of varying structural confidence by stratifying AlphaFold3 pLDDT scores at SNP and non-SNP positions (Figure 4A). Approximately 58% of non-synonymous SNPs fall in regions with high (70–90) or very high (>90) pLDDT scores, indicating that most structural predictions at variant sites can be interpreted with confidence. Non-synonymous SNPs nonetheless show a higher proportion of low-confidence positions than synonymous SNPs, which in turn show a higher proportion than non-SNP positions. Because low pLDDT scores correlate strongly with intrinsically disordered regions, this gradient reflects a relative enrichment of non-synonymous variants in disordered regions, consistent with the relaxed purifying selection previously reported for such regions in humans and yeast (Khan et al. 2015). To our knowledge, this is the first observation of this pattern at population scale in plants, and it illustrates the value of jointly analyzing structural and population level variant data.

**Figure 4.**
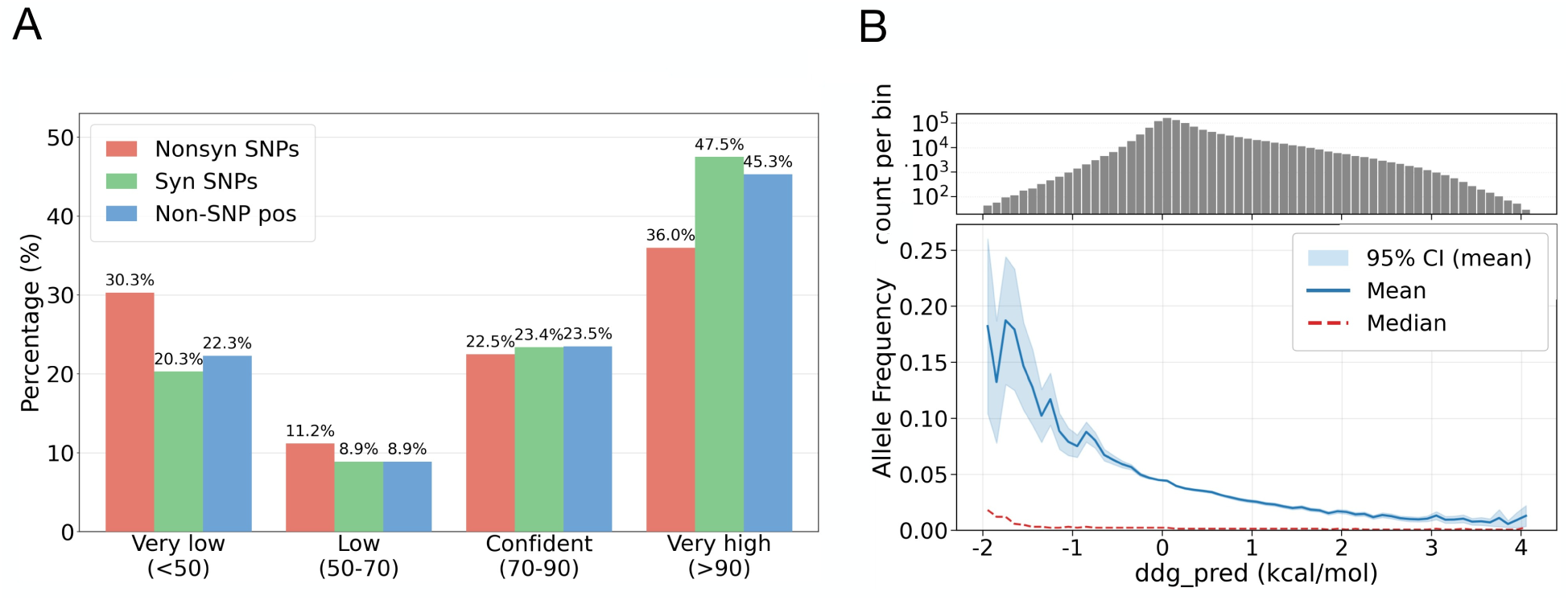
Global properties of the integrated SNPstar dataset. (A) Distribution of AlphaFold3 pLDDT confidence categories (very low, <50; low, 50–70; confident, 70–90; very high, >90) at non-synonymous SNP positions, synonymous SNP positions, and non-SNP positions in the reference proteotype structures. Non-synonymous SNPs are enriched in low-confidence regions and depleted in high-confidence regions relative to synonymous SNPs and non-SNP positions. Analysis was restricted to primary transcripts (.1 isoforms); positions carrying at least one non-synonymous SNP across the population were classified as non-synonymous. (B) Relationship between predicted thermodynamic stability change (ThermoMPNN ΔΔG, kcal mol⁻¹) and allele frequency across all variants. The upper panel shows the per-bin variant count (log scale); the lower panel shows the mean (solid line, with 95% confidence band) and median (dashed line) allele frequency as a function of predicted ΔΔG. SNPs with stabilizing predicted ΔΔG (negative values) occur at higher allele frequencies on average than destabilizing variants.

We next examined the relationship between predicted thermodynamic stability changes and allele frequency. Across all proteotypes, SNPs with stabilizing predicted ΔΔG values (negative ΔΔG) occur on average at significantly higher allele frequencies than destabilizing variants (Mann–Whitney U test, p < 0.001; Figure 4B). Although a stabilizing substitution does not necessarily improve protein function, this trend is consistent with purifying selection acting against destabilizing variants. A concrete example arises in the case study below: at HMA3 position 426, the substitution gaining a π–π stacking interaction is observed at substantially higher frequency than variants with a destabilizing amino acid change.

Together, these observations show what becomes possible when population-scale variation, structural confidence, and stability predictions are integrated in a single resource. The case studies below show how these structure-function relationships can be resolved for individual genes of interest.

## Case Study: *HMA3*

To illustrate how SNPstar is used in practice, we walk through a worked example with the gene *HEAVY METAL ATPASE 3* (*HMA3*; AT4G30120), encoding for a P-type ATPase known to limit long-distance transport of cadmium from root to shoot (Chao et al. 2012). *HMA3* is a useful test case because several of its natural variants have been functionally characterized in the literature, allowing us to compare SNPstar’s predictions with experimentally established outcomes. A step-by-step version of this walkthrough is available in the detailed tutorial on the SNPstar website.

### SNP overview and characterization

On the initial “Select transcripts and accessions” page, we enter the gene name *HMA3* (or its AGI ID, AT4G30120) and retain the default selection of all 1001 Genomes accessions. The interface then opens the ‘Results for selection’ view, where the sequence viewer (Figure 2C) provides a first overview of the SNPs in the gene. Several non-synonymous SNPs with high minor allele frequencies (≥ 10%) are immediately visible within the P-type heavy metal transporting ATPase domain of HMA3. We then switch to the SNP characterization table (Figure 3A) for a more detailed analysis.

The table lists a total of 176 SNPs in *HMA3*. Most of these are either synonymous, located in non-coding regions, or carry very low minor allele frequencies and are unlikely to be functionally consequential at the population level. For the remainder of this analysis we focus on non-synonymous SNPs with a minor allele frequency of at least 10%, a deliberately conservative cutoff that users can adjust depending on the gene of interest. Seven SNPs meet this threshold and are summarized in Figure 5.

**Figure 5.**
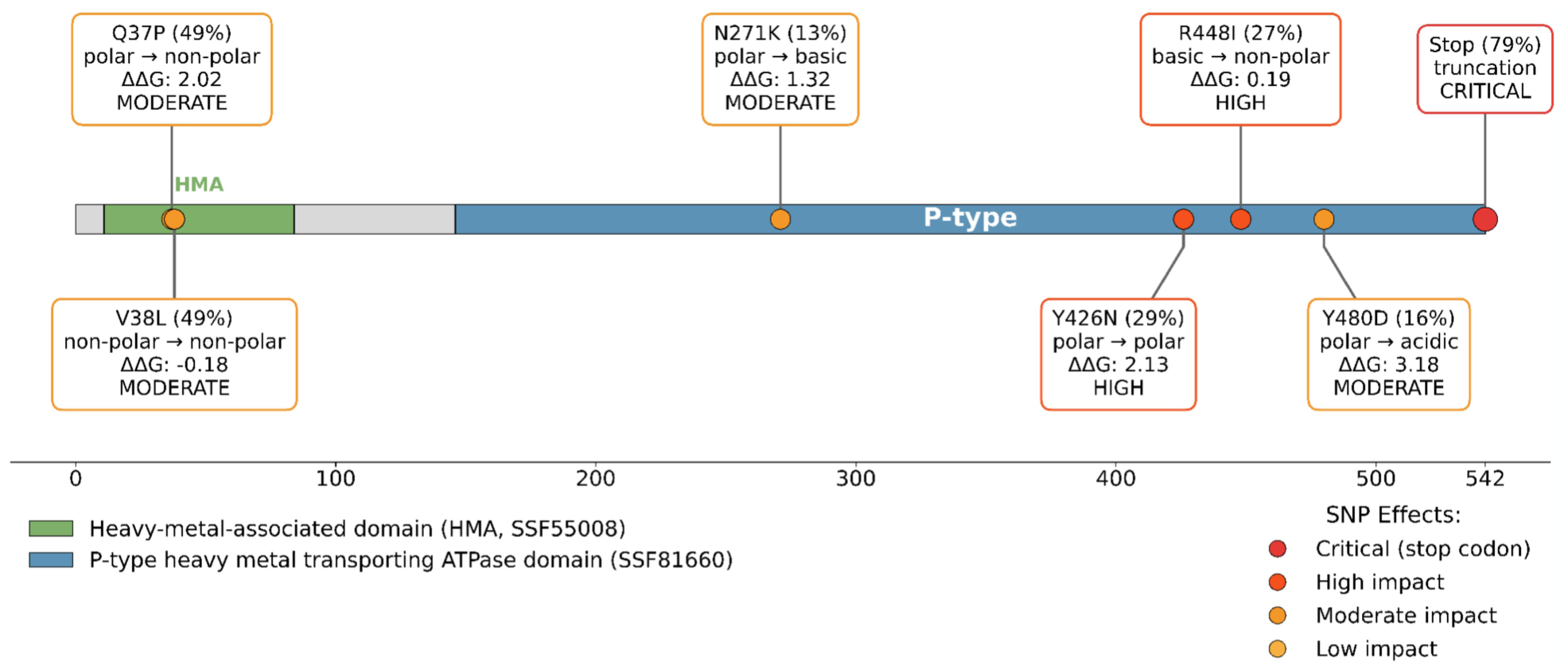
The seven HMA3 non-synonymous SNPs with minor allele frequency ≥ 10%, mapped onto the HMA3 domain architecture. Each variant is annotated with its position, allele frequency, change in chemical property class, FoldX-predicted ΔΔG, and SNPEff impact category. The N-terminal heavy-metal-associated (HMA, SSF55008) domain and the P-type heavy-metal-transporting ATPase domain (SSF81660) are shown in green and blue, respectively. The stop-codon variant at position 543 (79% frequency) represents stop-codon loss in non-Col-0 accessions, which produces an elongated functional protein relative to the truncated Col-0 reference. Visualization generated from SNPstar-exported data.

### Detailed SNP analysis

Six of the seven SNPs lie in the P-type heavy metal transporting ATPase domain, which spans most of the protein. Four of these (Q37P, N271K, Y426N, and R448I) show significant associations with leaf cadmium concentration in the AraGWAS catalog (Figure 5).

To assess potential functional impact, we combine the physicochemical property changes and predicted stability changes (FoldX and ThermoMPNN ΔΔG values; Dieckhaus et al. 2024) shown in SNPstar with detailed inspection of the proteotype structures. Where finer structural analysis was warranted - for example, identifying specific non-covalent interactions gained or lost - we exported the proteotype models from SNPstar and examined them with Arpeggio (Jubb et al. 2017) and PyMOL (Schrödinger, LLC 2015). Below we focus on three variants that illustrate the range of effects detectable through this combined approach: two missense substitutions (Y426N, Y480D) and one stop-codon variant (STOP543C) that, in non-Col-0 accessions, produces a longer protein than the Col-0 reference (Figure 6A, B).

**Figure 6.**
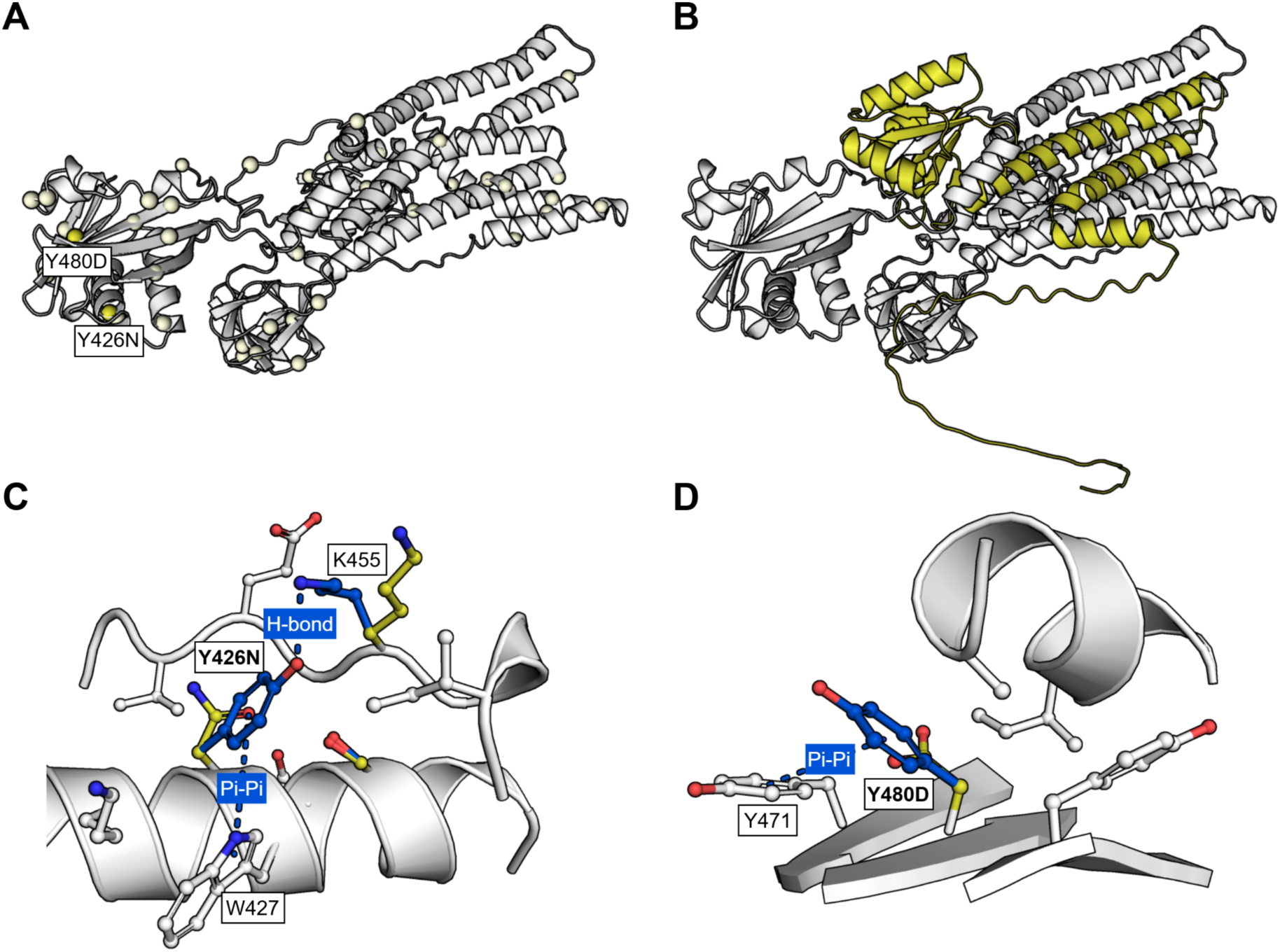
Structural visualization of selected HMA3 variants using AlphaFold3 models refined with FoldX. (A) Overview of the Col-0 HMA3 proteotype with non-synonymous SNP positions highlighted in yellow. (B) The stop-codon variant at position 543 yields a protein 218 amino acids longer than the Col-0 reference (additional residues shown in yellow; Col-0 reference in silver). (C) The Y426N substitution eliminates the hydrogen bond between Y426 and K455 and the π–π stacking interaction between Y426 and the adjacent W427. (D) The Y480D substitution removes the π–π stacking interaction with Y471 and the amide–π interactions with adjacent backbone atoms, contributing to the reduced thermodynamic stability predicted by FoldX and ThermoMPNN.

The Y426N substitution appears in 29% of accessions, suggesting selective advantages of either allele in specific environments, or alternatively genetic drift. It is located at the protein surface and disrupts multiple non-covalent interactions despite maintaining polarity. The mutation to asparagine eliminates both the hydrogen bond between Y426 and K455 and the edge-tilted π–π stacking interaction between Y426 and the adjacent W427. Arpeggio analysis (Figure 6C) shows that the asparagine replacement forms no compensatory hydrogen bonds, consistent with the positive ΔΔG values from FoldX and ThermoMPNN. Together, these features predict reduced thermodynamic stability of the variant. Experimental validation could proceed via thermal or chemical denaturation assays of recombinantly expressed Y426 and N426 variants.

The Y480D mutation replaces a surface-exposed tyrosine with aspartate, which is predicted to disrupt the π–π stacking interaction with Y471 as shown in Figure 6D. In line with this, FoldX and ThermoMPNN both predict reduced thermodynamic stability. Experimental validation has been provided by Chao et al. (2012), who showed that this mutation produces a loss-of-function phenotype of HMA3 with respect to cadmium transport, resulting in elevated cadmium concentrations in leaves.

The third variant we examined is a stop-codon mutation (STOP543C) that produces a protein 218 amino acids longer than the Col-0 reference (Figure 6B). The Col-0 truncated form is in fact the minor allele in this population, present in only 20% of accessions. Because SNPstar displays variants relative to the Col-0 reference, it correctly flags the stop-codon mutation but does not currently model the elongated protein sequence or its structural consequences, a limitation we discuss further below. SNPstar nonetheless detects the variant and assigns the affected accessions to a distinct haplotype and proteotype, allowing users to isolate them for downstream structural analysis with external tools. Chao et al. (2012) reported that the truncated Col-0 form, like the Y480D variant, is loss-of-function, with elevated cadmium concentrations in leaves.

Across these three variants, SNPstar surfaced candidates with strong predicted structural and functional consequences directly from the integrated dataset. For two of them, Y480D and the elongation variant, the predicted loss-of-function phenotype has already been experimentally established (Chao et al. 2012), demonstrating that SNPstar’s prioritization recovers known biology. The Y426N prediction remains to be tested experimentally and represents an example of a hypothesis a user could derive from SNPstar and validate in follow-up assays such as the thermal or chemical denaturation experiments outlined above.

As a final example, we used SNPstar to look for natural variation at residues whose functional importance has been established by site-directed mutagenesis. D397, the aspartic acid at position 397, is the canonical phosphorylation site in the catalytic cycle of P-type ATPases (Gravot et al. 2004). In that study, an EMS-induced D397A mutation, eliminating the carboxyl group required for phosphoryl transfer, abolished metal transport. The D397A substitution itself is not present in the 1001 Genomes dataset, but a different variant, D397E, does occur naturally at the same position.

Unlike the EMS-generated D397A mutation, D397E is a conservative substitution: glutamate retains the carboxyl group required for phosphoryl transfer, and both FoldX and ThermoMPNN predict minimal effects on overall stability. Glutamate’s side chain is, however, one methylene group longer and conformationally more flexible than aspartate’s side-chain. This geometric feature falls outside what ΔΔG predictors capture but could plausibly perturb the precise side-chain positioning required for phosphoryl transfer.

Because D397E targets the same catalytic residue as the experimentally characterized D397A mutation, one might speculate that the natural variant similarly affects HMA3 function, although the natural variant’s phenotype is, to our knowledge, unreported. It would therefore need to be tested directly, for example through functional assays of cadmium transport or phosphoproteomic analysis. Connecting population-level variants to mechanistically important residues in this way is a primary use case SNPstar is designed to support.

The HMA3 walkthrough illustrates the range of analyses SNPstar supports within a single interface: surfacing high-frequency non-synonymous variants, interpreting their predicted structural and stability consequences, comparing predictions against published experimental phenotypes, and generating testable hypotheses about variants whose function is not yet known. Two of the three high-frequency variants we examined (Y480D and the elongation variant) recapitulate known loss-of-function phenotypes; the third (Y426N) and the catalytic-residue variant D397E remain to be tested experimentally.

## Application: natural variation in *AGO2* slicer activity

The HMA3 case study recapitulated known biology; we next used SNPstar to address a previously uncharacterized molecular phenotype — natural variation in the slicer activity of ARGONAUTE 2 (*AGO2*). RNA interference is central to the plant antiviral response: ARGONAUTE (AGO) proteins load individual siRNA guide strands into the RNA-induced silencing complex (RISC), which base-pairs with complementary target RNAs and either cleaves them or represses their translation. Of the ten AGOs encoded by the *A. thaliana* genome, AGO1 and AGO2 act in antiviral defense, and loss of AGO2 aggravates disease symptoms in the *ago2-1* mutant (Harvey et al. 2011). AGO2 proteotypes have also been linked to viral susceptibility: accessions carrying the Col-0-like proteotype resist *Potato virus X*, whereas those with another (C24-like) proteotype are susceptible (Brosseau et al. 2020). Whether AGO2 proteotype diversity shapes responses to viruses that naturally infect Brassicaceae remains unknown, making biochemical characterization of catalytic (slicer) activity a promising route to connect genotype and phenotype.

AGO2 (encoded by *AT1G31280*) comprises an N-terminal intrinsically disordered region (N-IDR) with a glycine–arginine-rich (GR) motif, followed by the N-terminal, DUF1785, PAZ, MID and PIWI domains conserved across AGO proteins (Figure 7A). The N-terminal domain positions the guide–target duplex for cleavage and DUF1785 assists duplex unwinding; the MID and PAZ domains bind the 5′-monophosphate and the 3′ end of the guide strand, respectively, while the PIWI domain carries the catalytic triad that performs siRNA-guided target cleavage (Carbonell et al. 2012).

**Figure 7.**
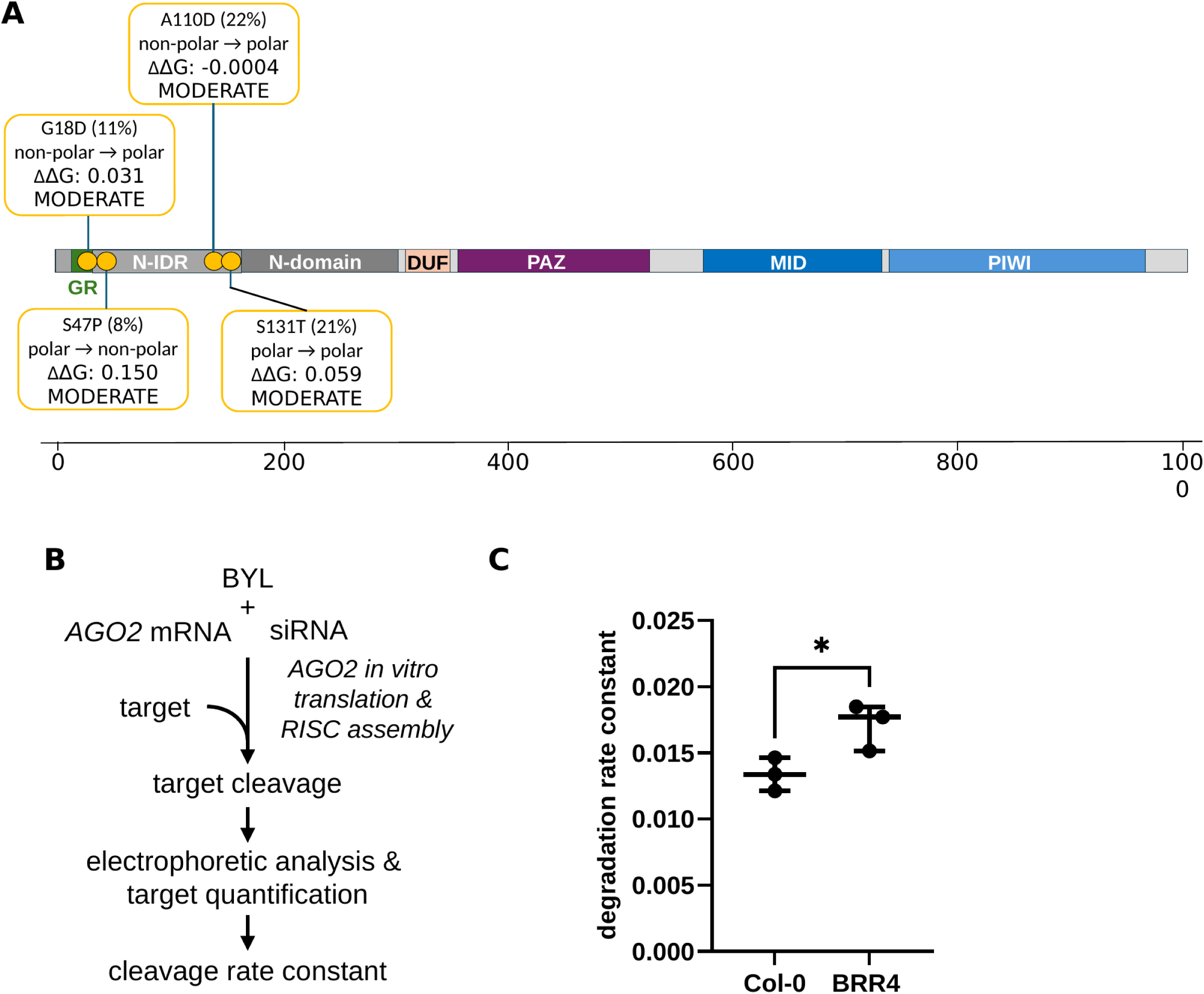
Effect of AGO2 proteoform variations on catalytic activity. (**A**) Schematic diagram of the AGO2 proteoform 3 (BRR4-AGO2) containing four amino acid substitutions derived from non-synonymous SNPs and a valine insertion. Conserved domains are indicated by boxes, and the glycine rich (GR) motif is highlighted in green. The amino acid changes are localized within the N-terminal disordered region (N-IDR). (**B**) Schematic illustration of the *in vitro* slicer assay. (**C**) Distribution of degradation rate constants (cleavage of target per second, measured as the reduction in target intensity over time) from individual kinetic assays, depicted as a box-and-whiskers plot (minimum to maximum, with all data points shown). Three independent experiments (see also Figure S2), statistical significance was assessed using an unpaired, two-tailed Student’s t-test: \**P* ≤ 0.05.

Using SNPstar, we catalogued 238 AGO2 proteotypes across the 1,135 sequenced *A. thaliana* accessions. Restricting to non-synonymous SNPs above 5% frequency, we identified 17 amino-acid positions that differ from the Col-0 reference, several co-selected within the same proteotype. Col-0 defines the most frequent proteotype (proteotype 1; 244 accessions). The third most frequent, proteotype 3 (88 accessions, 7.8%), carries four amino-acid changes in the N-IDR, present with and without an additional valine insertion (Figure 7A), and is found predominantly in accessions from the US.

To test whether this N-IDR variation affects catalytic activity, we cloned *AGO2* cDNAs from a representative proteotype-3 accession (BRR4) and from Col-0 and produced FLAG-tagged proteins by *in vitro* transcription and translation in the BYL system, a cytoplasmic extract of evacuolated *Nicotiana tabacum* BY-2 protoplasts. Slicer assays were performed in the same system (Figure 7B) as described (Gago-Zachert et al. 2019), using a target RNA of defined sequence; aliquots taken at successive time points were purified, resolved by denaturing gel electrophoresis, and quantified to derive a target-RNA degradation rate constant for each proteotype (Figure S2). AGO2^BRR4^ degraded the target significantly faster than AGO2^Col-0^ at 25 °C (Figure 7C). Because the BRR4 substitutions lie in the N-IDR, far from the catalytic PIWI domain, we hypothesize that this region acts indirectly rather than by reshaping the active site; for example by recruiting factors that enhance activity or by stabilizing the protein.

Notably, two of the four N-IDR substitutions carried by proteotype 3, A110D and S131T, occur at positions that Brosseau *et al*. (2020) previously identified as N-terminal polymorphisms associated with AGO2-dependent susceptibility to *Potato virus X*. That an unbiased, structure-aware screen in SNPstar converged on residues already shown by independent genetic and transgenic analysis to be functionally consequential provides further evidence that the tool prioritizes variants of genuine biological relevance, mirroring the *HMA3* result. The two studies are, however, complementary rather than redundant: Brosseau *et al*. assayed an *in planta* antiviral phenotype, whereas the slicer assay used here reports directly on the catalytic step, yielding a quantitative rate constant for target cleavage. The comparison also suggests that intrinsic slicer activity and antiviral protection are not simply equivalent, since AGO2^BRR4^ cleaves its target faster than AGO2^Col-0^ despite carrying the C24-type residues linked to viral susceptibility. A plausible reconciliation is that the N-IDR contributes to specific interactions with host and/or viral factors, rather than merely tuning catalysis, an interpretation that will require direct testing.

This application illustrates the full SNPstar workflow on a fresh biological question: proteotype grouping surfaced a high-frequency N-IDR variant set, geographic mapping tied it to a defined climatic origin, and the two contrasting accessions it nominated (Col-0 versus BRR4) translated directly into a measurable difference in slicer activity.

## Limitations

As the HMA3 case study made apparent, SNPstar has limitations that users should keep in mind.

On the data side, all analyses are anchored to the Col-0 reference genome; missing genotype calls in the 1001 Genomes data are imputed to the reference, which may underestimate true variation in those accessions. The current version is also restricted to *Arabidopsis thaliana* and to single-nucleotide variants: frameshifts, premature stop codons, altered start codons, and indels are detected and flagged in the SNP characterization table but are not yet incorporated into the structural models or proteotype definitions, so truncated or extended proteins must be analyzed with external tools (as illustrated by the elongation variant in the HMA3 case study). Combined effects of multiple SNPs within a single codon are reflected in the proteotype structures but not in the per-variant rows of the SNP characterization table.

On the prediction side, structures are computational rather than experimental (AlphaFold3 and FoldX), with accuracy varying across protein regions; the per-residue pLDDT scores in the interface should guide interpretation. To keep computation tractable and ensure adequate population representation, the structure viewer is also limited to proteotypes occurring above 1% allele frequency, which excludes rare variants that may nonetheless be functionally relevant in studies of local adaptation. Finally, the ΔΔG predictors capture thermodynamic effects but not chemistry-specific features such as catalytic geometry or PTM context; the D397E example in the case study shows how a variant with negligible predicted ΔΔG can still disrupt function. Low predicted ΔΔG should be read as structural tolerability, not as functional neutrality.

Extensions addressing several of these limitations - broader variant types, rare proteotypes, and additional species - are outlined in the Conclusions.

## Conclusions

SNPstar brings together population-scale variation, predicted protein structures, and stability estimates for *Arabidopsis thaliana* within a single gene-centric interface, allowing researchers to move from a list of SNPs to candidates for experimental validation without switching tools. By computing haplotypes and proteotypes alongside structural models, the resource enables analyses that span population frequency, sub-population structure, geographic distribution, and three-dimensional context; the scales at which non-synonymous variants are most informative. The HMA3 case study illustrates the practical payoff: starting from 176 raw variants, SNPstar surfaced a small set of high-confidence candidates, recapitulated two known loss-of-function alleles (the Y480D missense variant and the elongation variant; Chao et al. 2012), and generated a new structurally grounded hypothesis at a catalytic residue (D397E) for follow-up. The AGO2 application extends this in both directions: proteotype grouping nominated a high-frequency N-IDR variant set that independently converged on residues previously linked to antiviral function (Brosseau et al. 2020), while the two contrasting accessions it identified (Col-0 and BRR4) proved to differ measurably in *in vitro* slicer activity — a previously unquantified molecular phenotype. Together, the two case studies show SNPstar recovering established biology and generating new mechanistic hypotheses from the same integrated dataset. Planned developments include expansion to long-read pan-genome assemblies (Lian et al. 2024), broader variant coverage (CNVs, indels, structural variants), more complete structural annotations, on-the-fly analysis of user-defined variants, and protein complex and interaction models. The underlying architecture is designed to be extensible to other species, opening a path toward comparative analyses across the plant kingdom. Lastly, a potentially powerful add-on would be a SNP score that predicts potential functional consequences of SNPs based on genomic, structural, and phenotypic data catalogued in SNPstar.

By making the joint analysis of natural variation and predicted protein structure tractable for non-specialist users, SNPstar lowers a long-standing barrier between population genomics and structural biology; a foundation we expect to be useful well beyond the Arabidopsis community.

## Author contributions

BS ran the genomic analyses with support of FP, programed SNPstar and wrote the manuscript with MQ; LP and GK computed the protein structures with contributions of CT and PLK; SB and JT tested SNPstar and developed the manual; MT and JG maintain the SNPstar webtool; SGZ and SEB conceptualized and performed the functional studies with the AGO2 proteoforms; IG, PLK and MQ conceptually developed the project.

## Acknowledgements

Martin Luther University core funding for enabling us to initiate the development of SNPstar. We thank Lasse Feldhahn for his support in the early establishment phase. Most of all, we thank all SNP2Prot members (https://snp2prot.uni-halle.de/) and many colleagues not named individually for beta-testing SNPstar and providing valuable feedback.

## Supplementary material

Table S1. Comparison of functionality across different variant analysis webtools for *Arabidopsis thaliana*.

Figure S1. Basic step-by-step tutorial introducing SNPstar’s features, accessible via the *Getting Started* link on the SNPstar homepage.

Figure S2. Single-turnover cleavage kinetics of Col-0 and BRR4 AGO2 in three independent replicates.

## Funding

This research was supported by the Deutsche Forschungsgemeinschaft (DFG, project 514901783) through the Collaborative Research Centre 1664 “Plant Proteoform Diversity – SNP2Prot” (projects D01 and D02). SGZ and SEB were funded by the Deutsche Forschungsgemeinschaft (project BE 1885/15-1).

## Conflict of interest statement

None declared.

## Notes

### Competing Interest Statement

The authors have declared no competing interest.

