## Supplemental Table 1 for "SNPstar: A Web Server Linking Allelic Variation to Protein Structure and Function in *Arabidopsis thaliana*"

| Functionality \ Webtool | 1001 genomes tools | Plants Ensembl | UniProt | Variant viewer | Viva webtool | SNPstar |
|---|---|---|---|---|---|---|
| **Online availability** | ✗ | ✔ | ✔ | ✔ | ✗ | ✔ |
| **SNP visualization within isoforms** | ✗ | ✔ | ✔ | ✗ | ✔ | ✔ |
| **SNP characterization & functional analysis** | ✔ | ✔ | ✗ | ✗ | ✔ | ✔ |
| **Association of SNPs with accessions** | ✔ | ✗ | ✗ | ✗ | ✔ | ✔ |
| **Custom accession subset selection** | ✔ | ✗ | ✗ | ✗ | ✔ | ✔ |
| **Haplotype/proteotype computation** | ✗ | ✗ | ✗ | ✗ | ✗ | ✔ |
| **Batch analysis of multiple isoforms** | ✗ | ✗ | ✗ | ✗ | ✗ | ✔ |
| **GWAS matrix generation** | ✗ | ✗ | ✗ | ✗ | ✗ | ✔ |
| **Protein structures with integrated SNPs** | ✗ | ✗ | ✔ | ✗ | ✗ | ✔ |

**Table S1.** Comparison of functionality across different variant analysis webtools for *Arabidopsis thaliana*. The tools compared are: 1001 genomes tools (1001 Genomes Project 2024b), Plants Ensembl (Yates et al. 2021b), UniProt (Bateman et al. 2022), Variant viewer ("Variant Viewer" 2024), and Viva webtool (Hamm et al. 2019). Green checkmarks indicate supported functionality, red crosses indicate unsupported. SNPstar provides comprehensive functionality across all capabilities, while other tools are more limited in their feature sets.
