## Supplemental Figure 1 for "SNPstar: A Web Server Linking Allelic Variation to Protein Structure and Function in *Arabidopsis thaliana*"

### SNPstar - Getting started in 33 steps

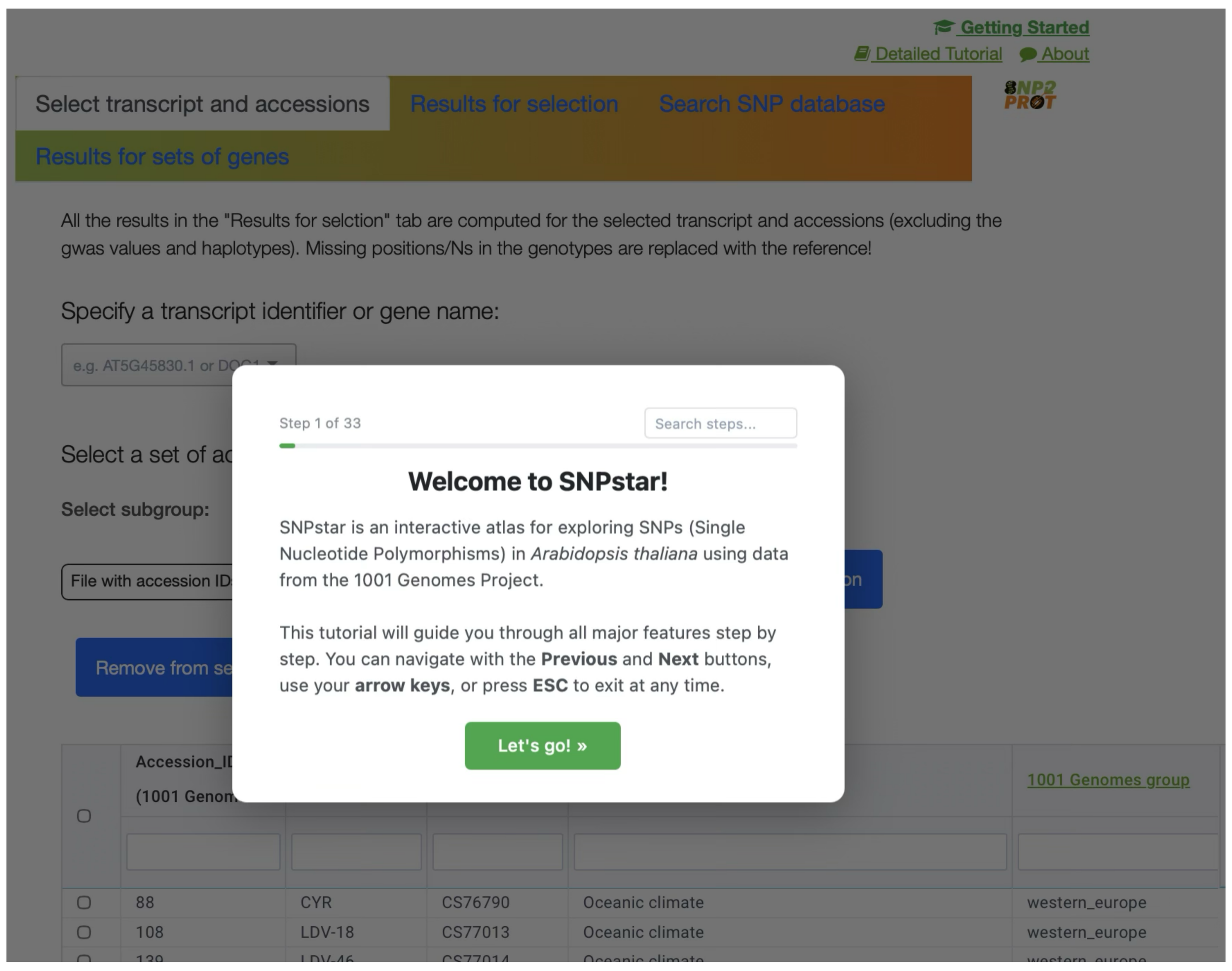


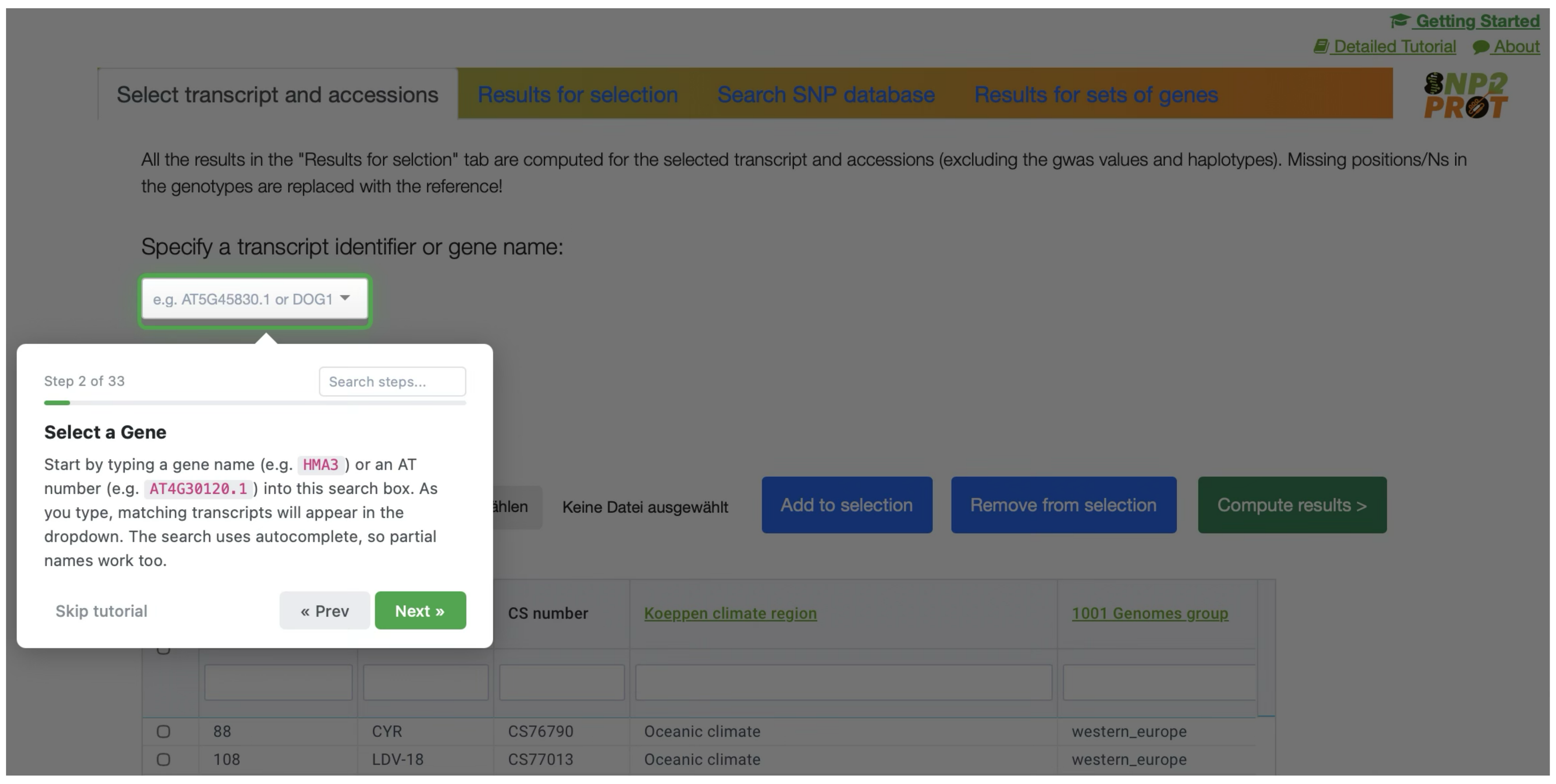


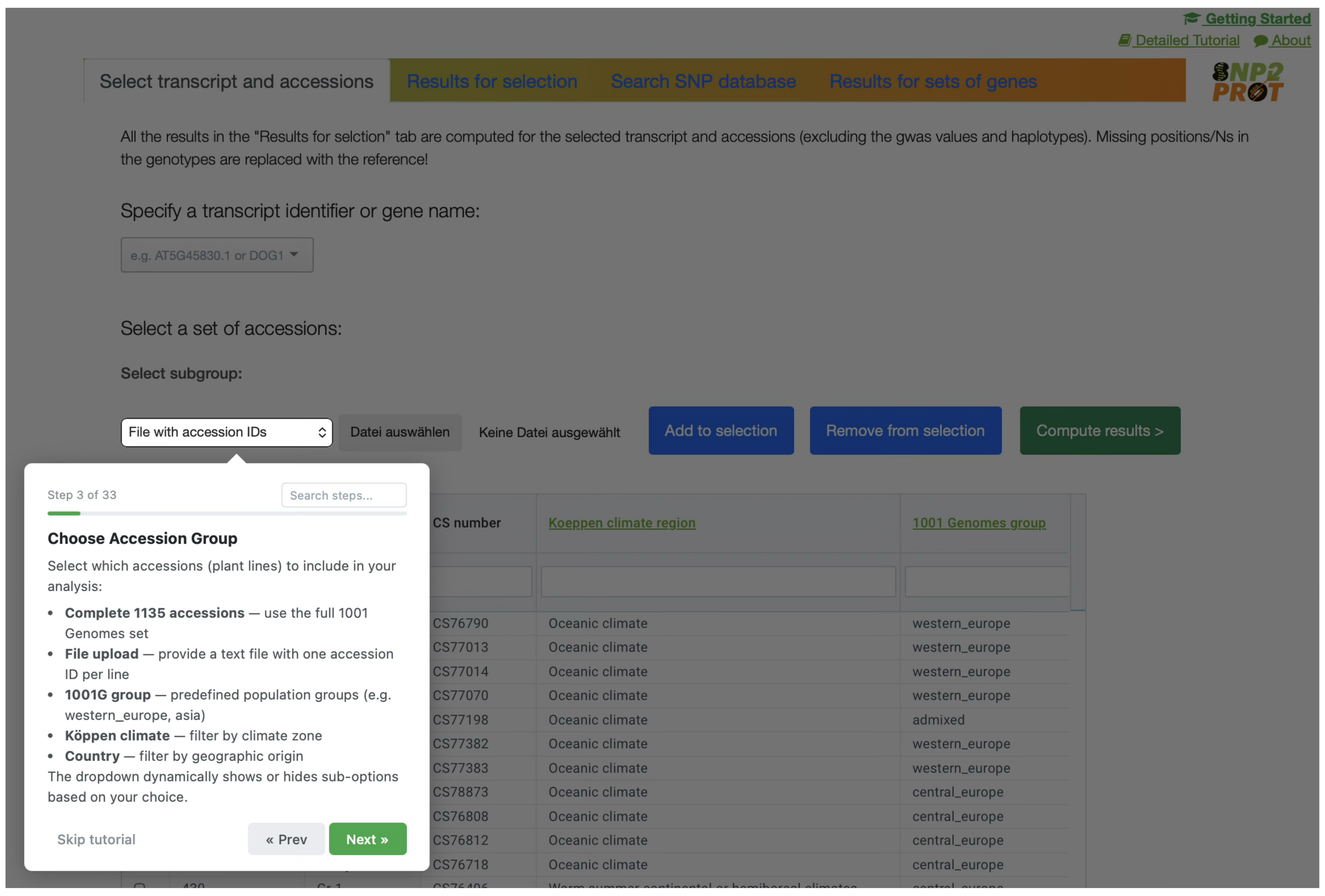


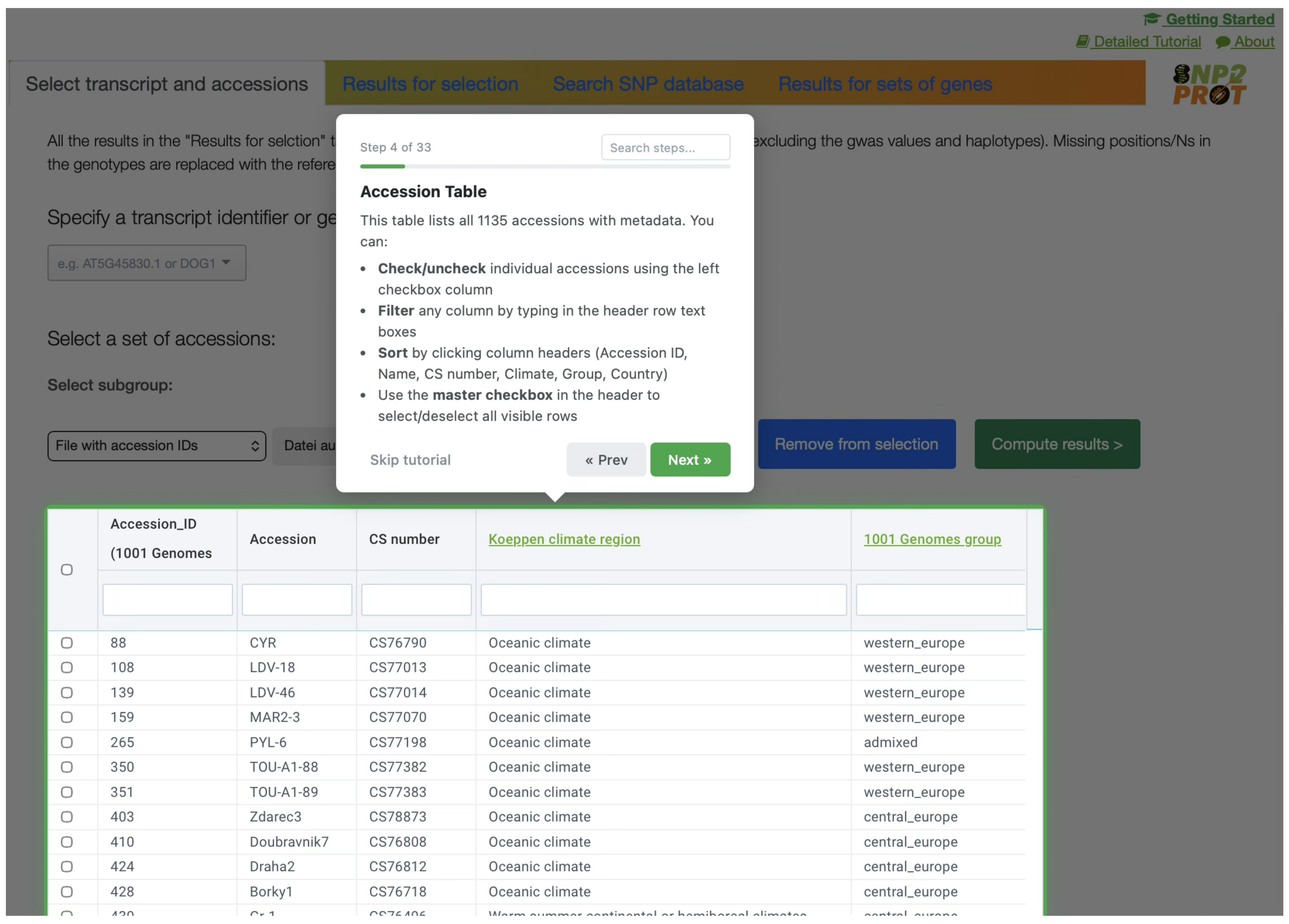


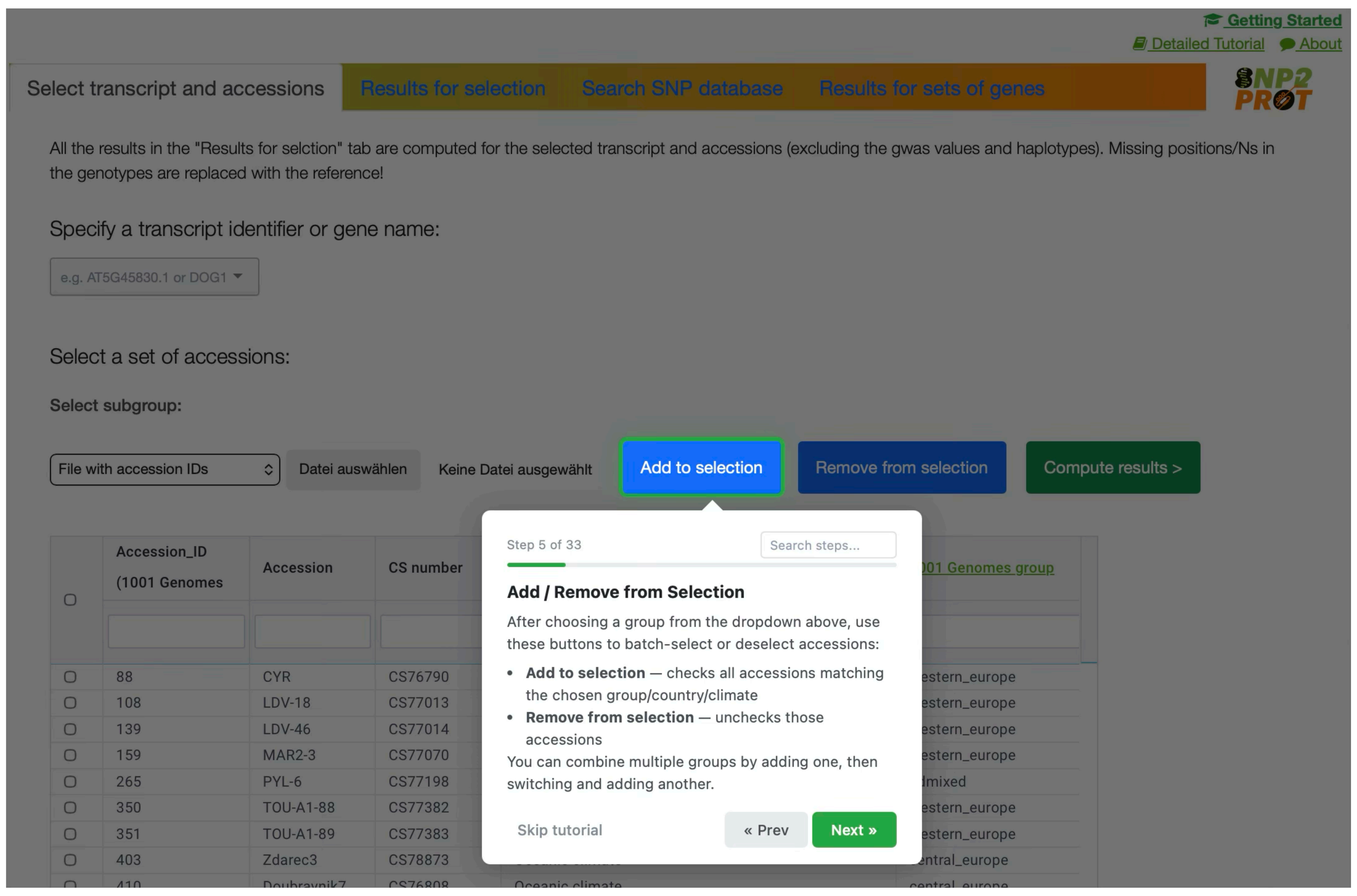


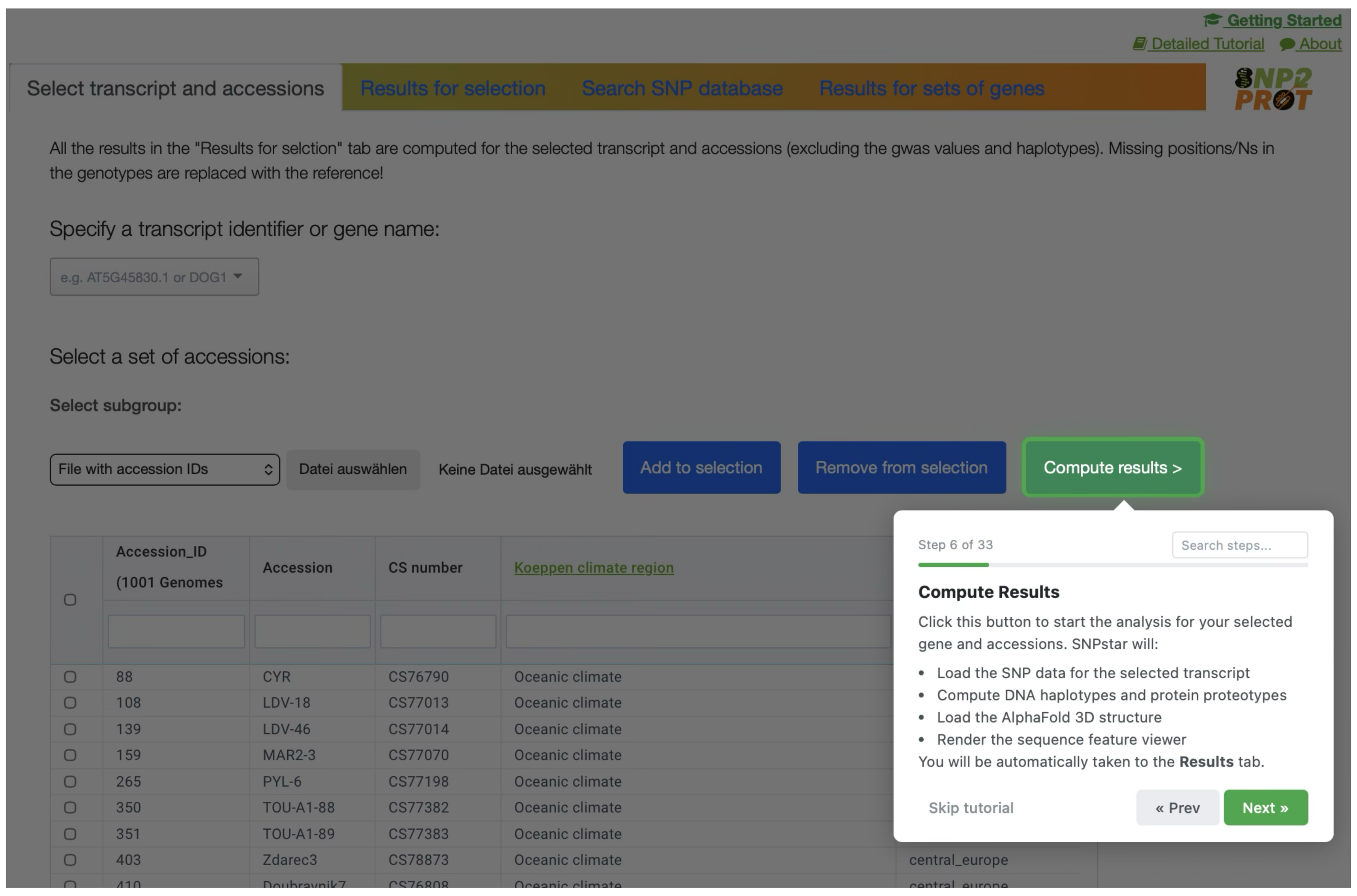

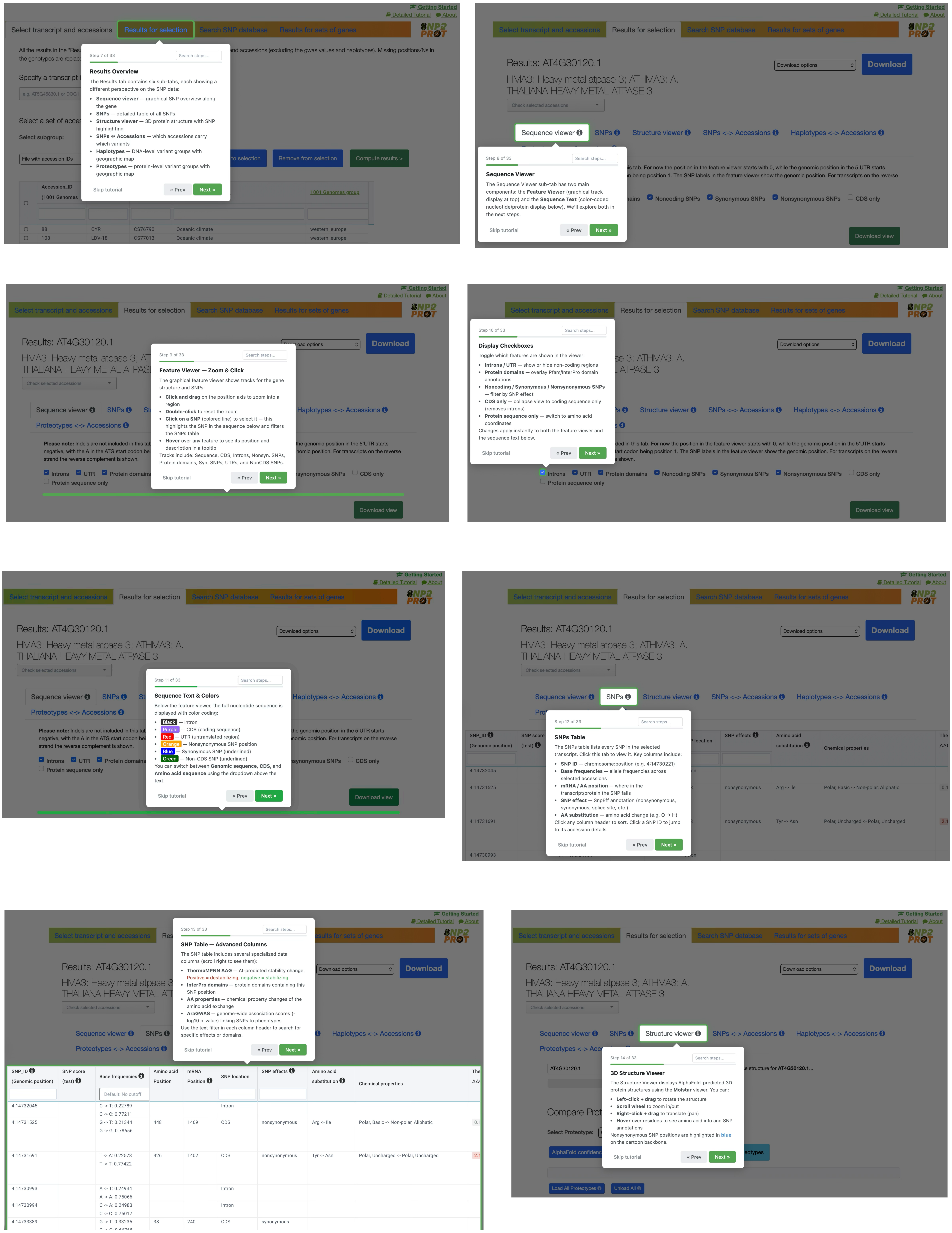
Getting Started
Detailed Tutorial
About
Select transcript and accessions
Results for selection
Search SNP database
Results for sets of genes
Step 7 of 33
Results Overview
The Results tab contains six sub-tabs, each showing a different perspective on the SNP data:
Sequence viewer — graphical SNP overview along the gene
SNPs — detailed table of all SNPs
Structure viewer — 3D protein structure with SNP highlighting
SNPs ↔ Accessions — which accessions carry which variants
Haplotypes — DNA-level variant groups with geographic map
Proteotypes — protein-level variant groups with geographic map
Skip tutorial
« Prev
Next »
Results: AT4G30120.1
HMA3: Heavy metal atpase 3; ATHMA3: A. THALIANA HEAVY METAL ATPASE 3
Download options
Download
Check selected accessions
Sequence viewer
SNPs
Structure viewer
SNPs <-> Accessions
Haplotypes <-> Accessions
Proteotypes <-> Accessions
Step 8 of 33
Sequence Viewer
The Sequence Viewer sub-tab has two main components: the Feature Viewer (graphical track display at top) and the Sequence Text (color-coded nucleotide/protein display below). We'll explore both in the next steps.
Download view
Step 9 of 33
Feature Viewer — Zoom & Click
The graphical feature viewer shows tracks for the gene structure and SNPs:
Click and drag on the position axis to zoom into a region
Double-click to reset the zoom
Click on a SNP (colored line) to select it — this highlights the SNP in the sequence below and filters the SNPs table
Hover over any feature to see its position and description in a tooltip
Tracks include: Sequence, CDS, Introns, Nonsyn. SNPs, Protein domains, Syn. SNPs, UTRs, and NonCDS SNPs.
Please note: Indels are not included in this tab. For now the position in the feature viewer starts with 0, while the genomic position in the 5'UTR starts with the A in the ATG start codon being position 1. The SNP labels in the feature viewer show the genomic position. For transcripts on the reverse strand the reverse complement is shown.
Introns
UTR
Protein domains
Noncoding SNPs
Synonymous SNPs
Nonsynonymous SNPs
CDS only
Protein sequence only
Step 10 of 33
Display Checkboxes
Toggle which features are shown in the viewer:
Introns / UTR — show or hide non-coding regions
Protein domains — overlay Pfam/InterPro domain annotations
Noncoding / Synonymous / Nonsynonymous SNPs — filter by SNP effect
CDS only — collapse view to coding sequence only (removes introns)
Protein sequence only — switch to amino acid coordinates
Changes apply instantly to both the feature viewer and the sequence text below.
Step 11 of 33
Sequence Text & Colors
Below the feature viewer, the full nucleotide sequence is displayed with color coding:
Black — Intron
Purple — CDS (coding sequence)
Red — UTR (untranslated region)
Orange — Nonsynonymous SNP position
Blue — Synonymous SNP (underlined)
Green — Non-CDS SNP (underlined)
You can switch between Genomic sequence, CDS, and Amino acid sequence using the dropdown above the text.
Step 12 of 33
SNPs Table
The SNPs table lists every SNP in the selected transcript. Click this tab to view it. Key columns include:
SNP ID — chromosome:position (e.g. 4:14730221)
Base frequencies — allele frequencies across selected accessions
mRNA / AA position — where in the transcript/protein the SNP falls
SNP effect — SnpEff annotation (nonsynonymous, synonymous, splice site, etc.)
AA substitution — amino acid change (e.g. Q → H)
Click any column header to sort. Click a SNP ID to jump to its accession details.
Step 13 of 33
SNP Table — Advanced Columns
The SNP table includes several specialized data columns (scroll right to see them):
ThermoMPNN ΔΔG — AI-predicted stability change. Positive = destabilizing, negative = stabilizing
InterPro domains — protein domains containing this SNP position
AA properties — chemical property changes of the amino acid exchange
AraGWAS — genome-wide association scores (-log10 p-value) linking SNPs to phenotypes
Use the text filter in each column header to search for specific effects or domains.
SNP_ID (Genomic position)
SNP score (test)
Base frequencies
Amino acid Position
mRNA Position
SNP location
SNP effects
Amino acid substitution
Chemical properties
Default: No cutoff
4:14732045 C -> T: 0.22789 C -> C: 0.77211 Intron
4:14731525 G -> T: 0.21344 G -> G: 0.78656 448 1469 CDS nonsynonymous Arg -> Ile Polar, Basic -> Non-polar, Aliphatic
4:14731691 T -> A: 0.22578 T -> T: 0.77422 426 1402 CDS nonsynonymous Tyr -> Asn Polar, Uncharged -> Polar, Uncharged
4:14730993 A -> T: 0.24934 A -> A: 0.75066 Intron
4:14730994 C -> A: 0.24983 C -> C: 0.75017 Intron
4:14733389 G -> T: 0.33235 38 240 CDS synonymous
Step 14 of 33
3D Structure Viewer
The Structure Viewer displays AlphaFold-predicted 3D protein structures using the Molstar viewer. You can:
Left-click + drag to rotate the structure
Scroll wheel to zoom in/out
Right-click + drag to translate (pan)
Hover over residues to see amino acid info and SNP annotations
Nonsynonymous SNP positions are highlighted in blue on the cartoon backbone.
Compare Proteotypes
Select Proteotype:
AlphaFold confidence
Load All Proteotypes
Unload All

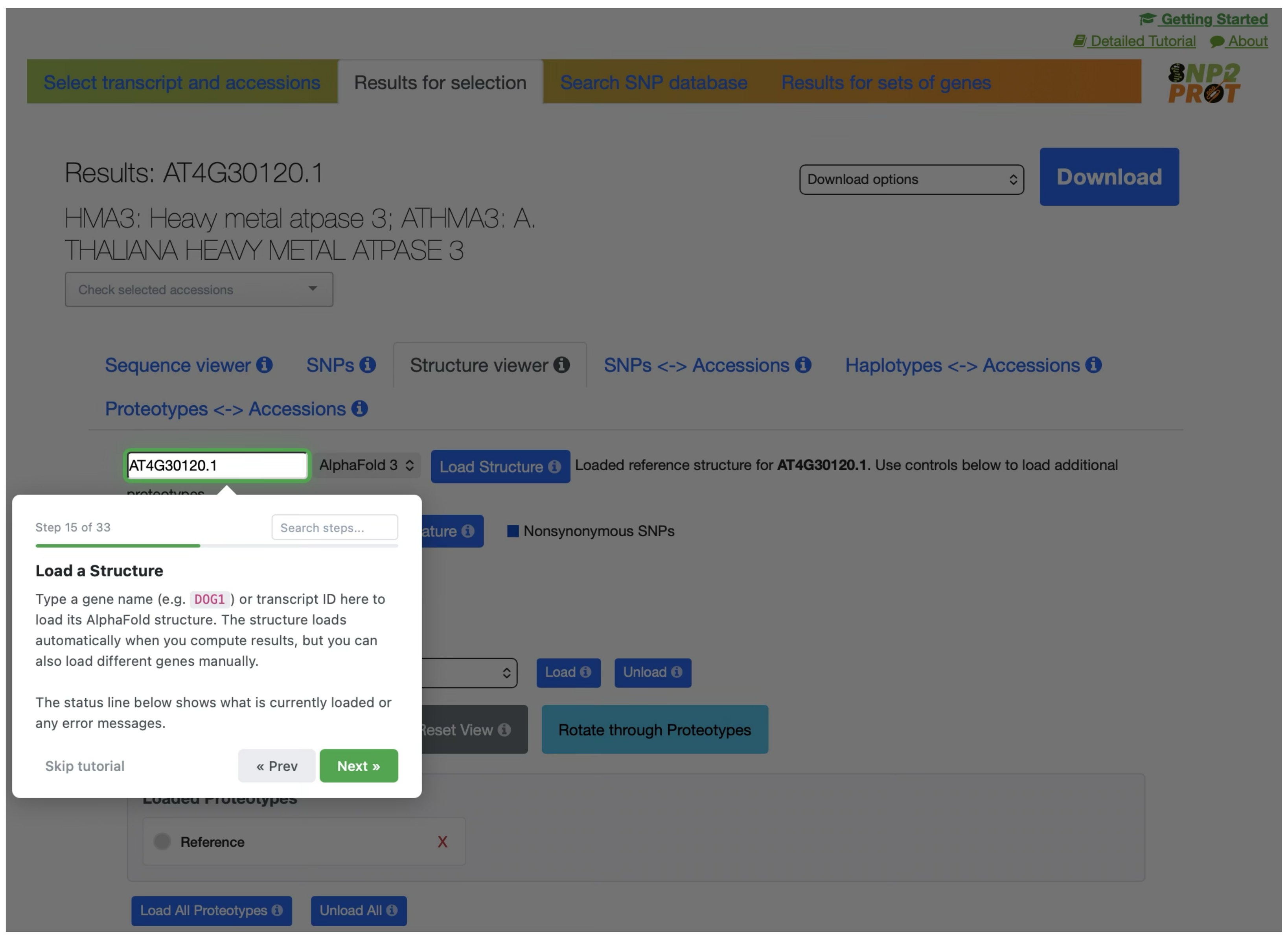
Step 15 of 33
Load a Structure
Type a gene name (e.g. DOG1) or transcript ID here to load its AlphaFold structure. The structure loads automatically when you compute results, but you can also load different genes manually.
The status line below shows what is currently loaded or any error messages.
Skip tutorial
« Prev
Next »

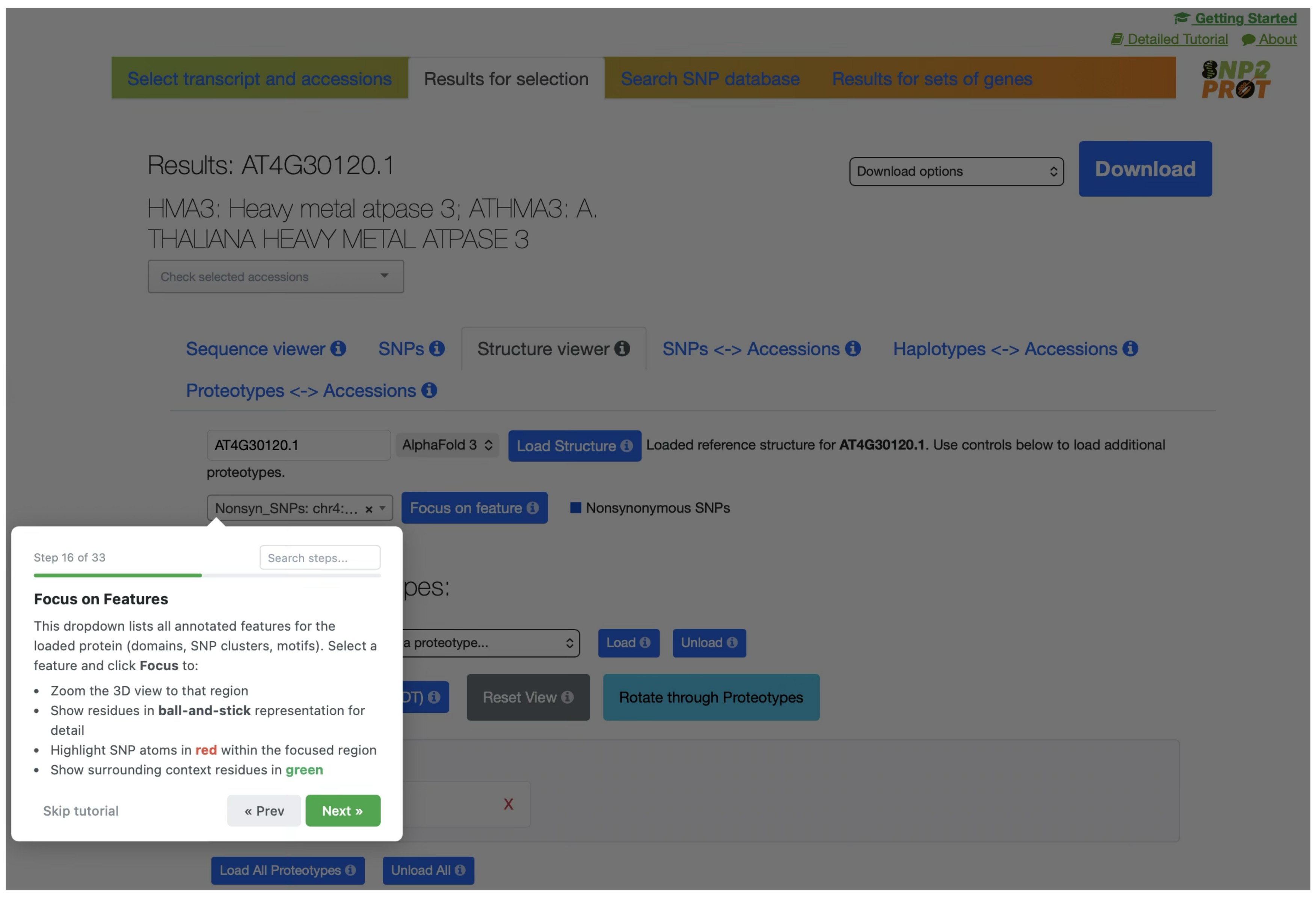
Step 16 of 33
Focus on Features
This dropdown lists all annotated features for the loaded protein (domains, SNP clusters, motifs). Select a feature and click Focus to:
Zoom the 3D view to that region
Show residues in ball-and-stick representation for detail
Highlight SNP atoms in red within the focused region
Show surrounding context residues in green
Skip tutorial
« Prev
Next »

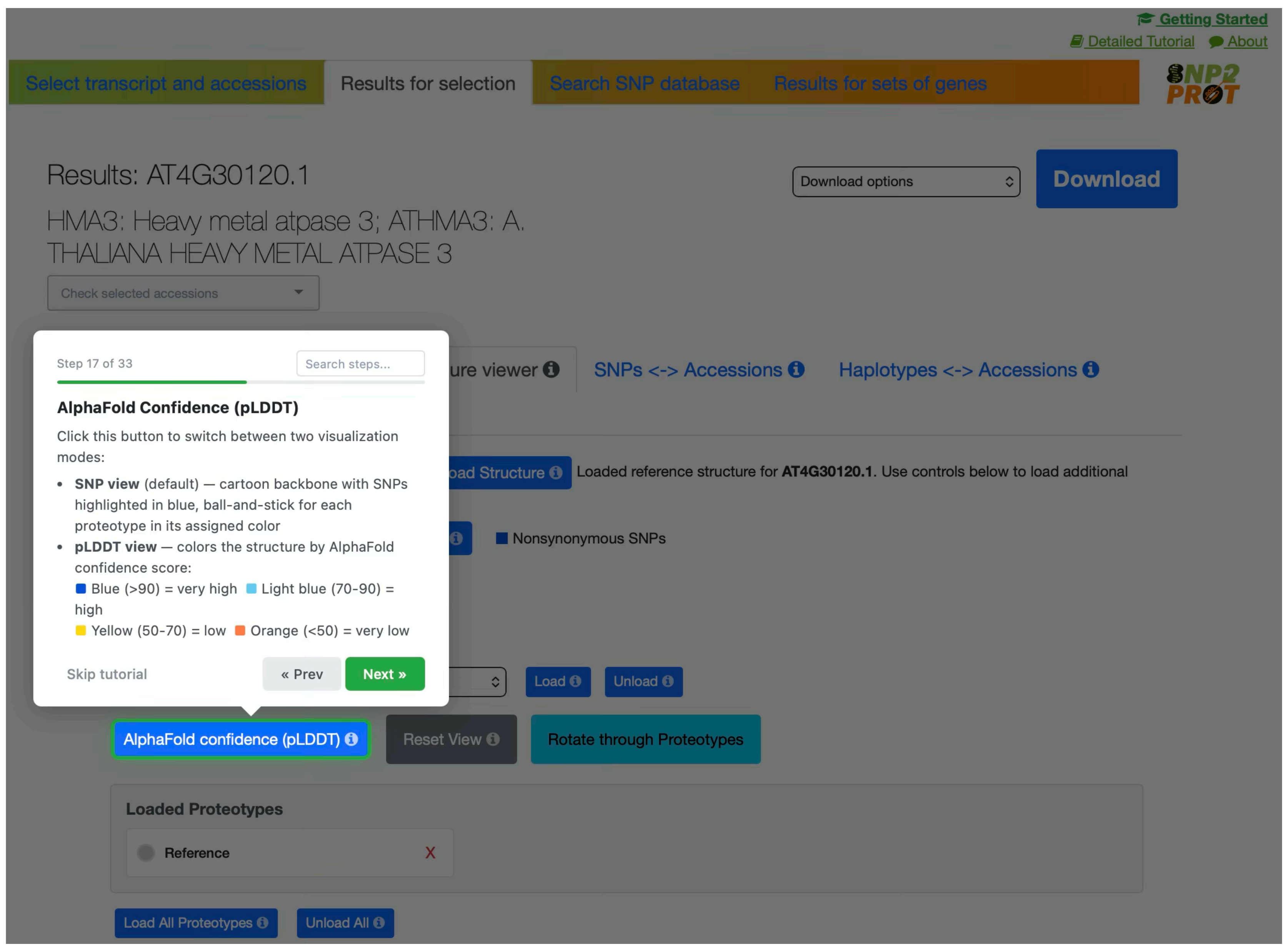
Step 17 of 33
AlphaFold Confidence (pLDDT)
Click this button to switch between two visualization modes:
SNP view (default) — cartoon backbone with SNPs highlighted in blue, ball-and-stick for each proteotype in its assigned color
pLDDT view — colors the structure by AlphaFold confidence score:
Blue (>90) = very high Light blue (70-90) = high
Yellow (50-70) = low Orange (<50) = very low
Skip tutorial
« Prev
Next »
AlphaFold confidence (pLDDT)

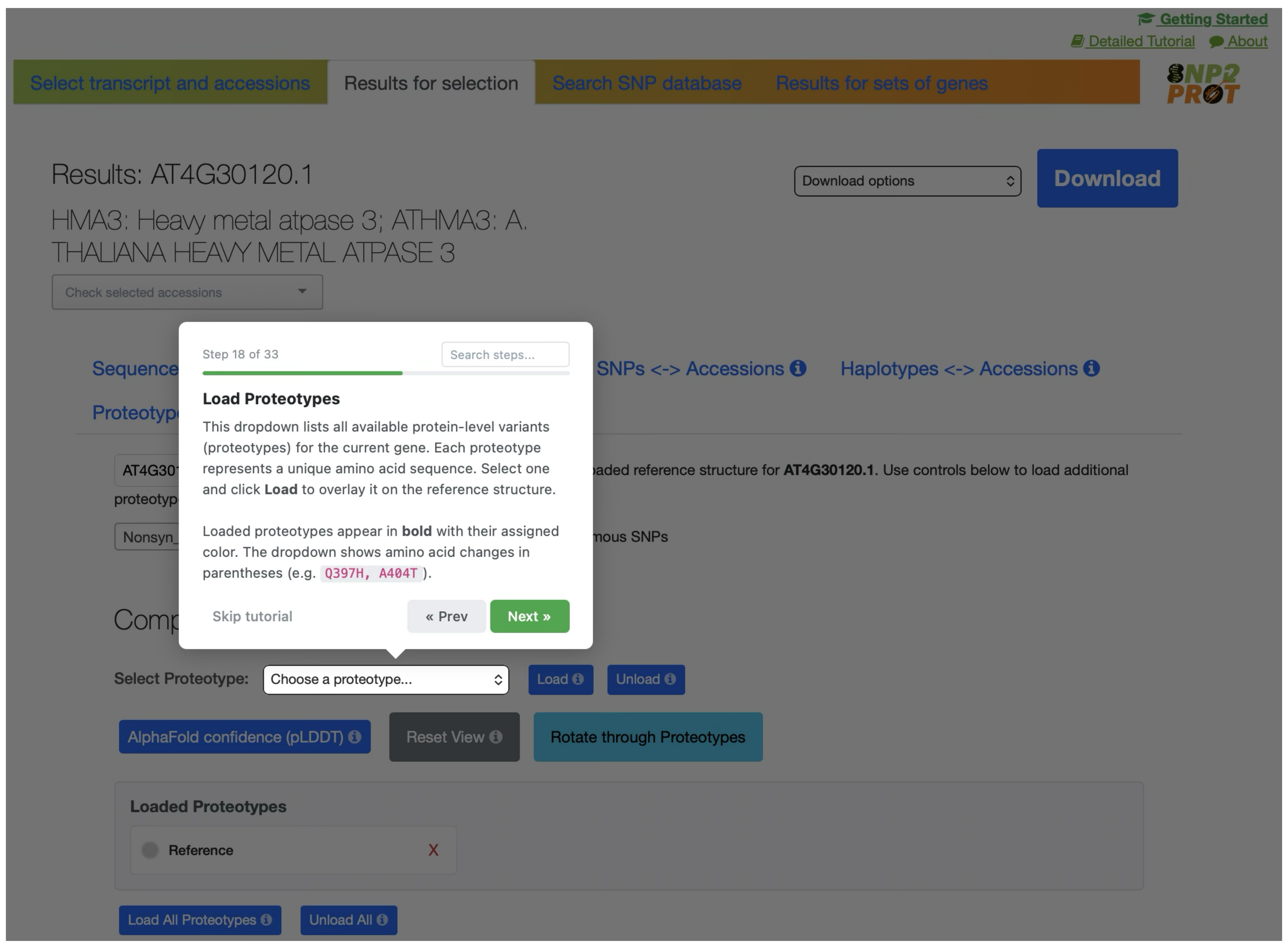
Step 18 of 33
Load Proteotypes
This dropdown lists all available protein-level variants (proteotypes) for the current gene. Each proteotype represents a unique amino acid sequence. Select one and click Load to overlay it on the reference structure.
Loaded proteotypes appear in bold with their assigned color. The dropdown shows amino acid changes in parentheses (e.g. Q397H, A404T).
Skip tutorial
« Prev
Next »
Select Proteotype:
Choose a proteotype...

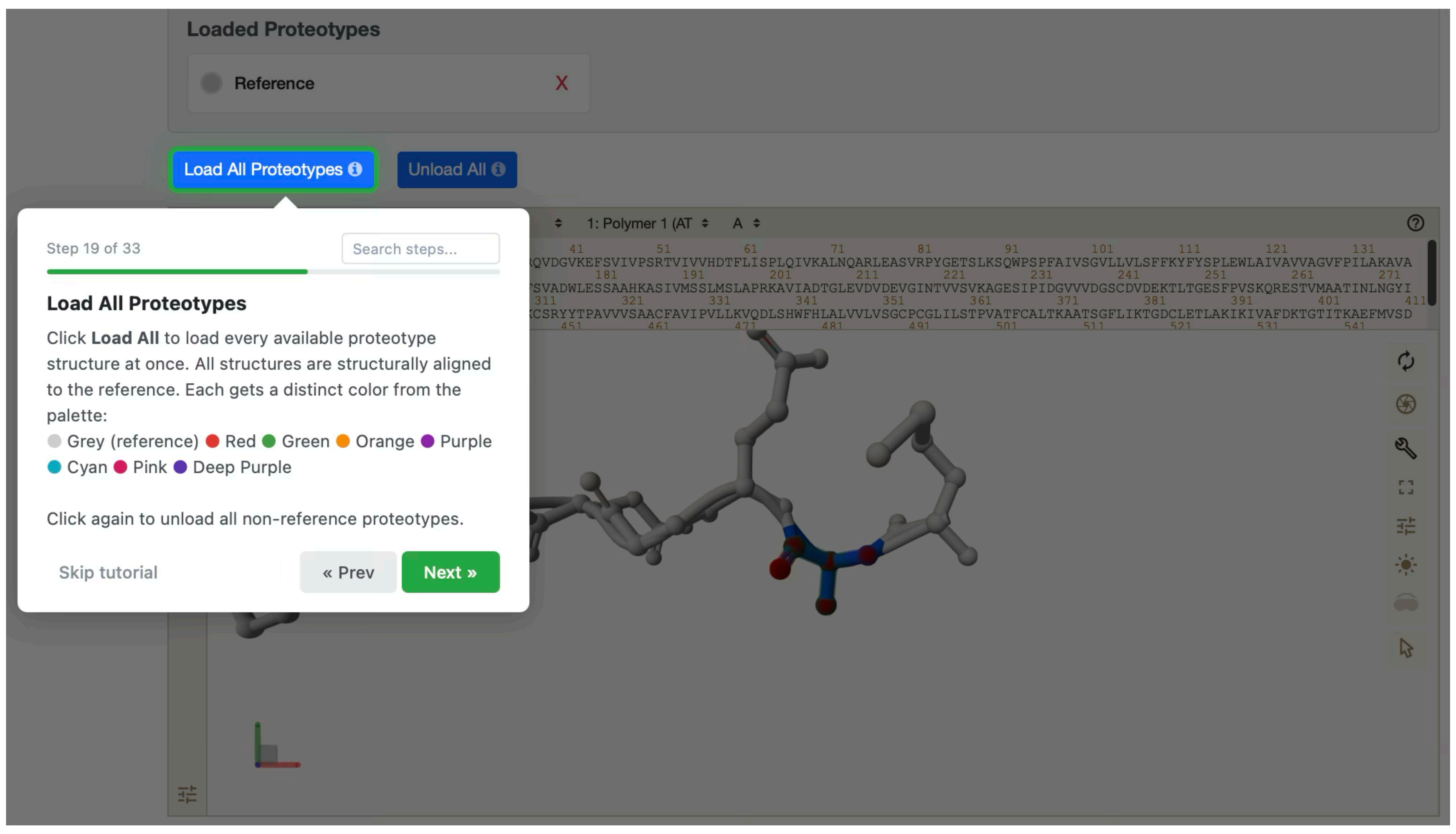
Loaded Proteotypes
Reference
Load All Proteotypes
Unload All
Step 19 of 33
Load All Proteotypes
Click Load All to load every available proteotype structure at once. All structures are structurally aligned to the reference. Each gets a distinct color from the palette:
Grey (reference) Red Green Orange Purple Cyan Pink Deep Purple
Click again to unload all non-reference proteotypes.
Skip tutorial
« Prev
Next »

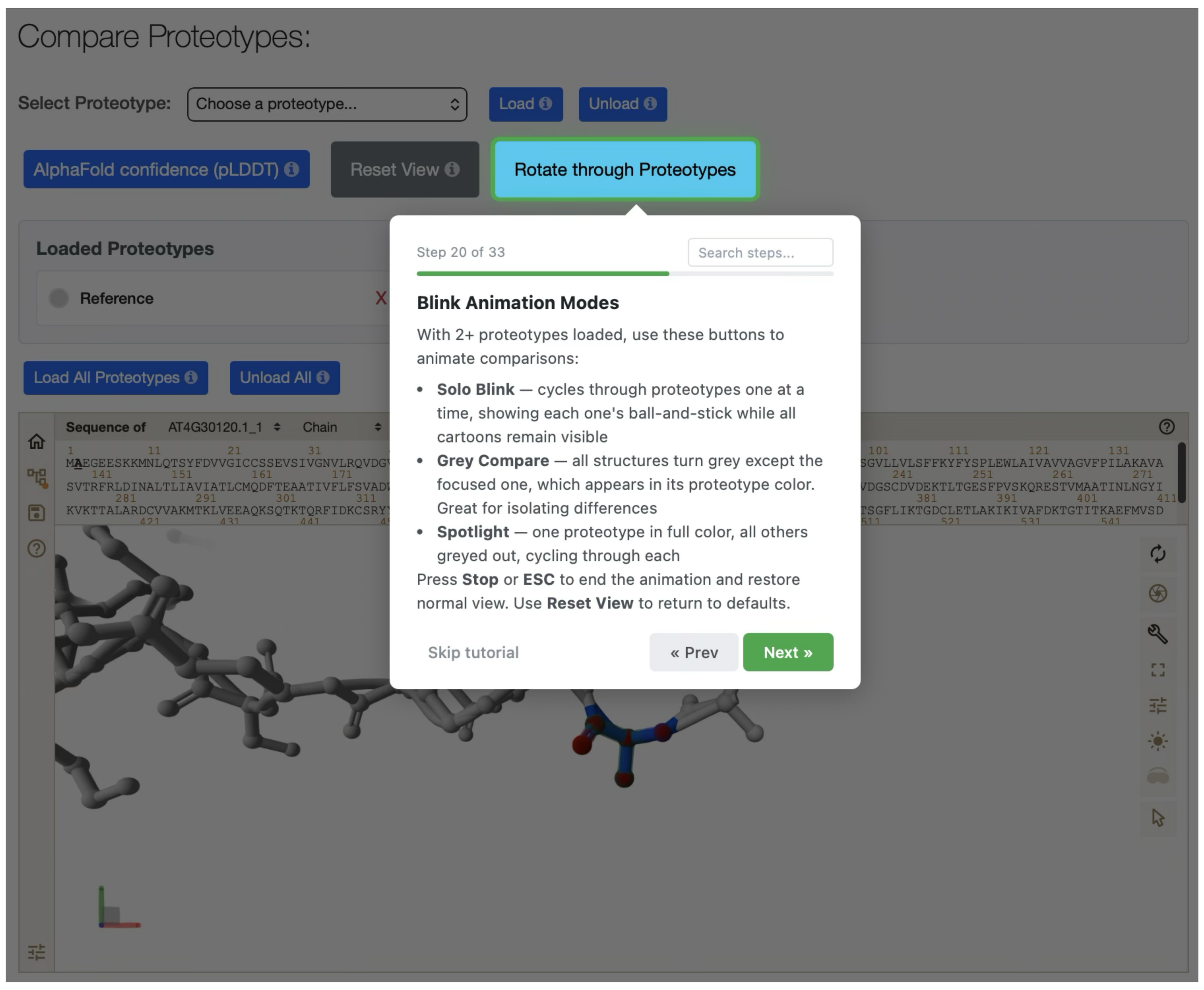
Compare Proteotypes:
Select Proteotype:
Choose a proteotype...
Load
Unload
AlphaFold confidence (pLDDT)
Reset View
Rotate through Proteotypes
Loaded Proteotypes
Reference
Load All Proteotypes
Unload All
Step 20 of 33
Blink Animation Modes
With 2+ proteotypes loaded, use these buttons to animate comparisons:
Solo Blink — cycles through proteotypes one at a time, showing each one's ball-and-stick while all cartoons remain visible
Grey Compare — all structures turn grey except the focused one, which appears in its proteotype color. Great for isolating differences
Spotlight — one proteotype in full color, all others greyed out, cycling through each
Press Stop or ESC to end the animation and restore normal view. Use Reset View to return to defaults.
Skip tutorial
« Prev
Next »

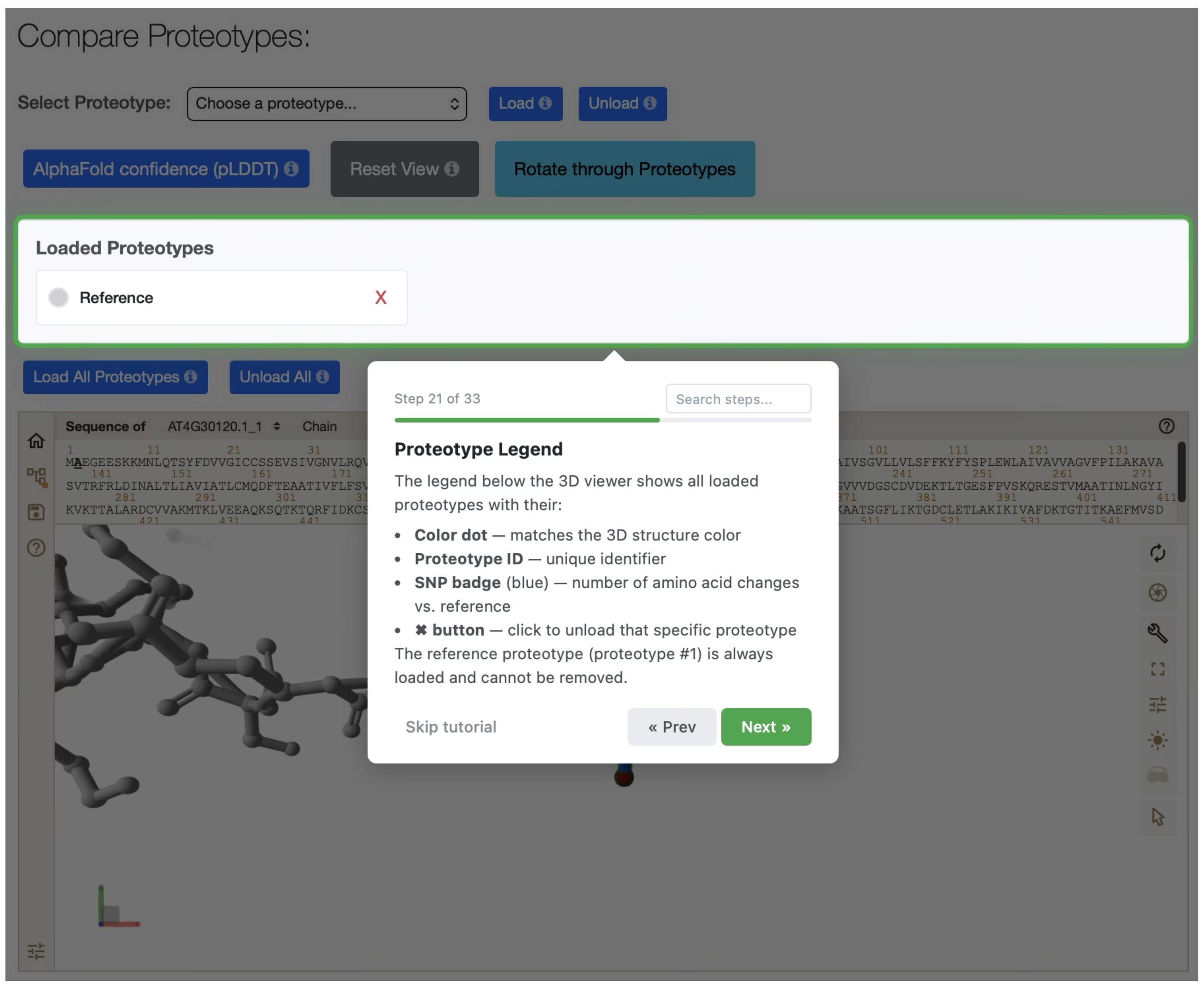

Compare Proteotypes:
Select Proteotype:
Choose a proteotype...
Load
Unload
AlphaFold confidence (pLDDT)
Reset View
Rotate through Proteotypes
Loaded Proteotypes
Reference
Load All Proteotypes
Unload All
Step 21 of 33
Search steps...
Proteotype Legend
The legend below the 3D viewer shows all loaded proteotypes with their:
• Color dot — matches the 3D structure color
• Proteotype ID — unique identifier
• SNP badge (blue) — number of amino acid changes vs. reference
• ✖ button — click to unload that specific proteotype
The reference proteotype (proteotype #1) is always loaded and cannot be removed.
Skip tutorial
« Prev
Next »


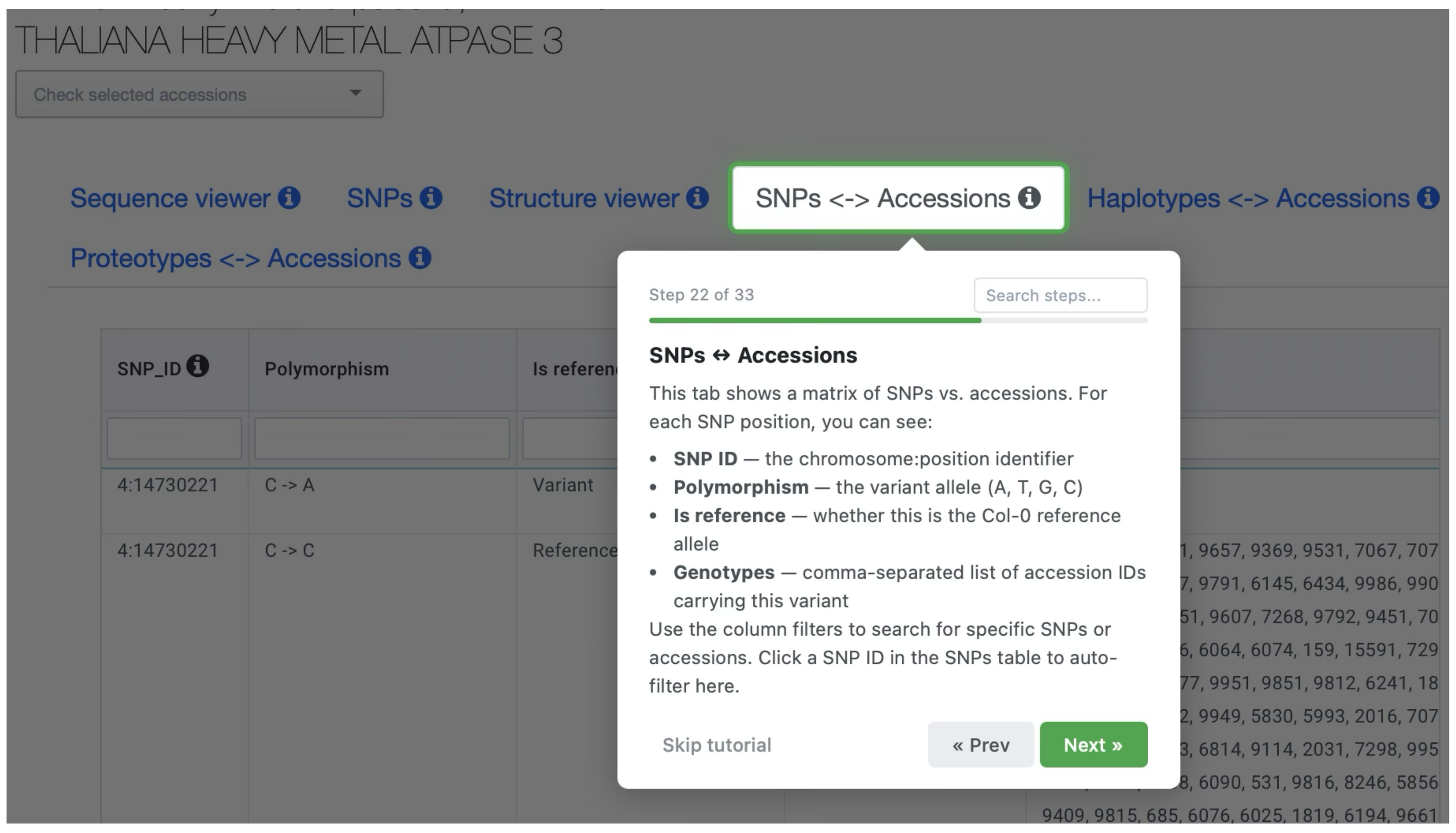

THALIANA HEAVY METAL ATPASE 3
Check selected accessions
Sequence viewer
SNPs
Structure viewer
SNPs <-> Accessions
Haplotypes <-> Accessions
Proteotypes <-> Accessions
SNP_ID
Polymorphism
4:14730221
C -> A
Variant
4:14730221
C -> C
Reference
Step 22 of 33
Search steps...
SNPs ↔ Accessions
This tab shows a matrix of SNPs vs. accessions. For each SNP position, you can see:
• SNP ID — the chromosome:position identifier
• Polymorphism — the variant allele (A, T, G, C)
• Is reference — whether this is the Col-0 reference allele
• Genotypes — comma-separated list of accession IDs carrying this variant
Use the column filters to search for specific SNPs or accessions. Click a SNP ID in the SNPs table to auto-filter here.
Skip tutorial
« Prev
Next »


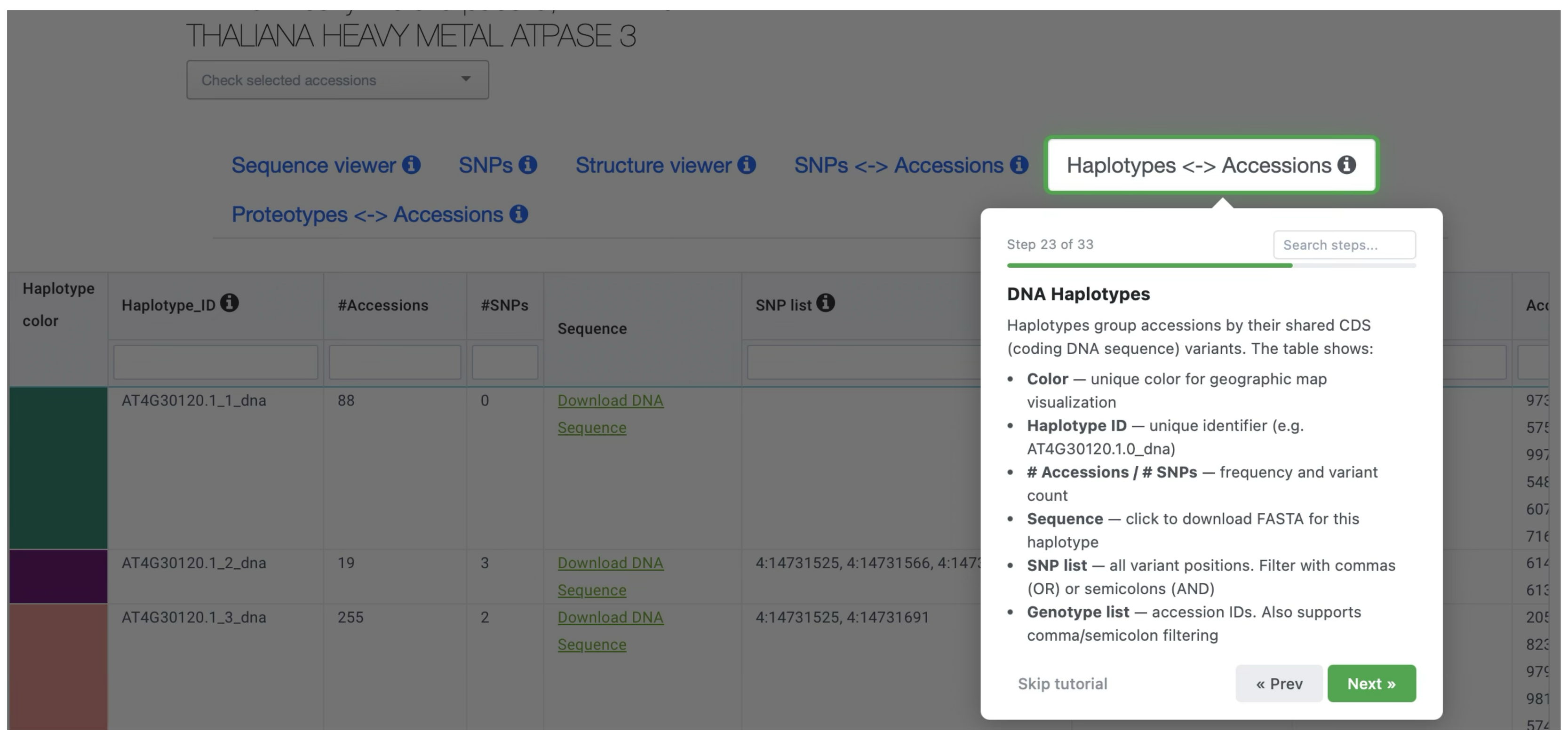

THALIANA HEAVY METAL ATPASE 3
Sequence viewer
SNPs
Structure viewer
SNPs <-> Accessions
Haplotypes <-> Accessions
Proteotypes <-> Accessions
Haplotype color
Haplotype_ID
#Accessions
#SNPs
Sequence
SNP list
AT4G30120.1_1_dna
88
0
Download DNA Sequence
AT4G30120.1_2_dna
19
3
AT4G30120.1_3_dna
255
2
Step 23 of 33
Search steps...
DNA Haplotypes
Haplotypes group accessions by their shared CDS (coding DNA sequence) variants. The table shows:
• Color — unique color for geographic map visualization
• Haplotype ID — unique identifier (e.g. AT4G30120.1.0_dna)
• # Accessions / # SNPs — frequency and variant count
• Sequence — click to download FASTA for this haplotype
• SNP list — all variant positions. Filter with commas (OR) or semicolons (AND)
• Genotype list — accession IDs. Also supports comma/semicolon filtering
Skip tutorial
« Prev
Next »


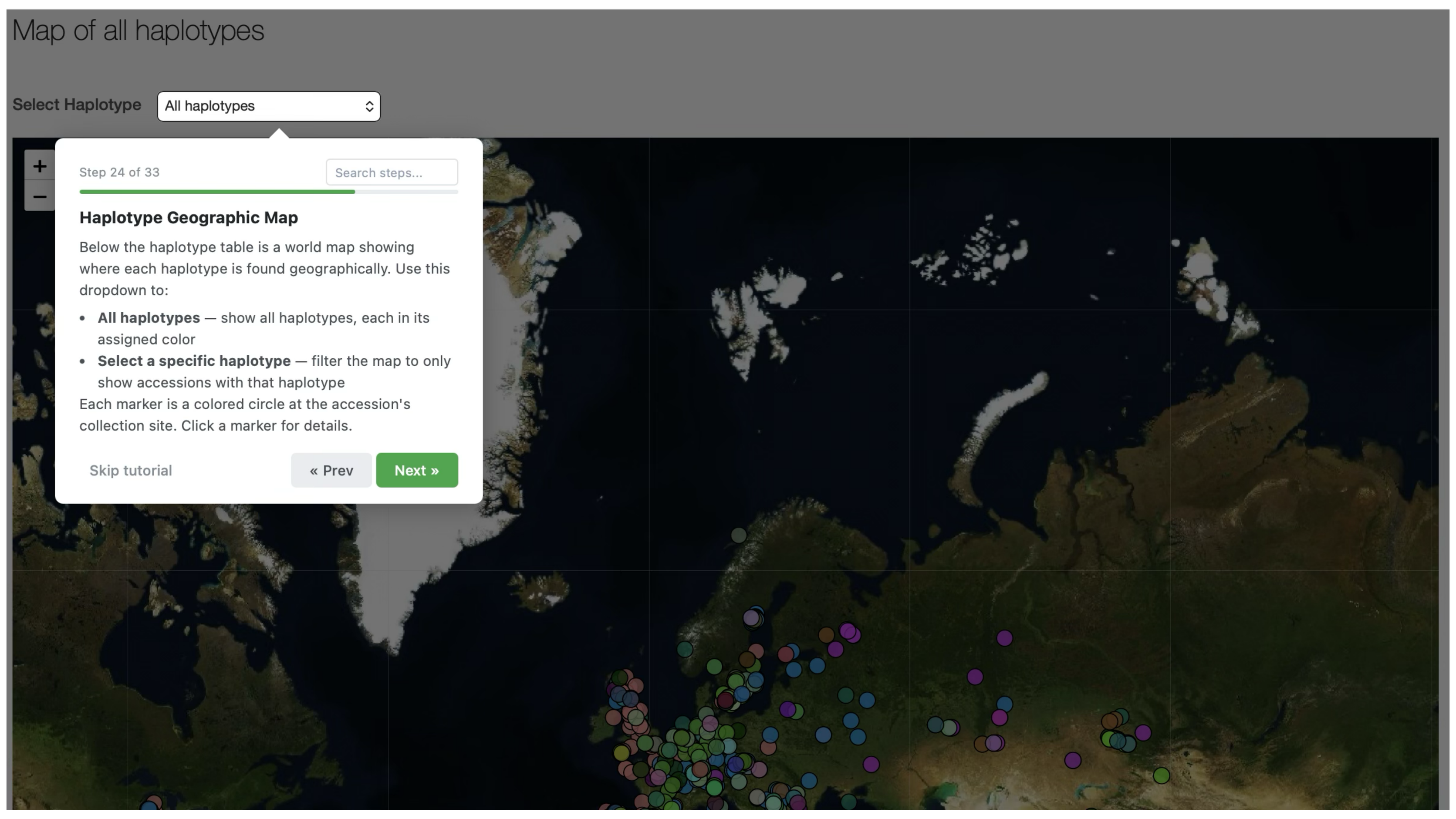

Map of all haplotypes
Select Haplotype
All haplotypes
Step 24 of 33
Search steps...
Haplotype Geographic Map
Below the haplotype table is a world map showing where each haplotype is found geographically. Use this dropdown to:
• All haplotypes — show all haplotypes, each in its assigned color
• Select a specific haplotype — filter the map to only show accessions with that haplotype
Each marker is a colored circle at the accession's collection site. Click a marker for details.
Skip tutorial
« Prev
Next »


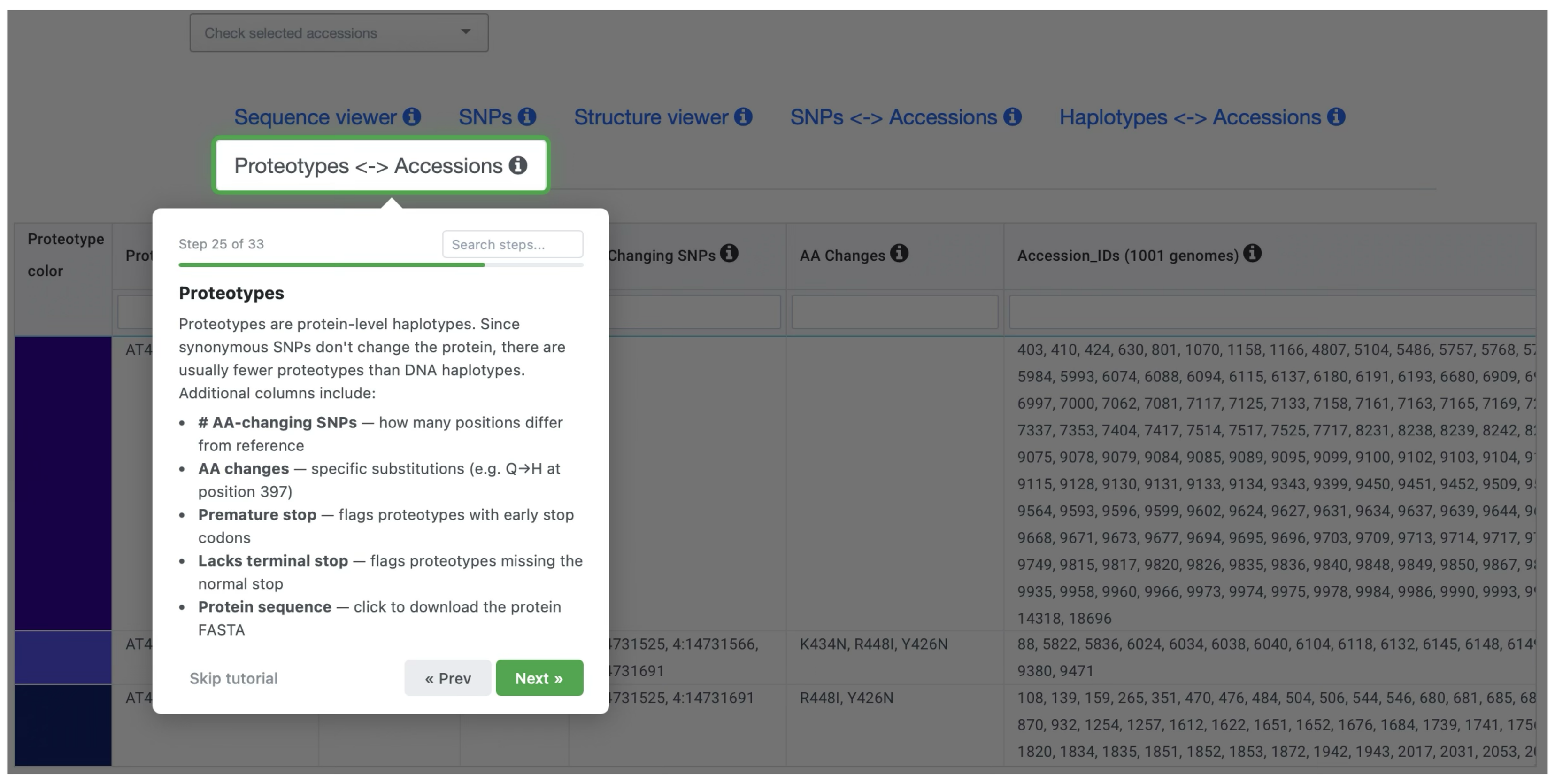

Check selected accessions
Sequence viewer
SNPs
Structure viewer
SNPs <-> Accessions
Haplotypes <-> Accessions
Proteotypes <-> Accessions
AA Changes
Accession_IDs (1001 genomes)
K434N, R448I, Y426N
R448I, Y426N
Step 25 of 33
Search steps...
Proteotypes
Proteotypes are protein-level haplotypes. Since synonymous SNPs don't change the protein, there are usually fewer proteotypes than DNA haplotypes. Additional columns include:
• # AA-changing SNPs — how many positions differ from reference
• AA changes — specific substitutions (e.g. Q→H at position 397)
• Premature stop — flags proteotypes with early stop codons
• Lacks terminal stop — flags proteotypes missing the normal stop
• Protein sequence — click to download the protein FASTA
Skip tutorial
« Prev
Next »


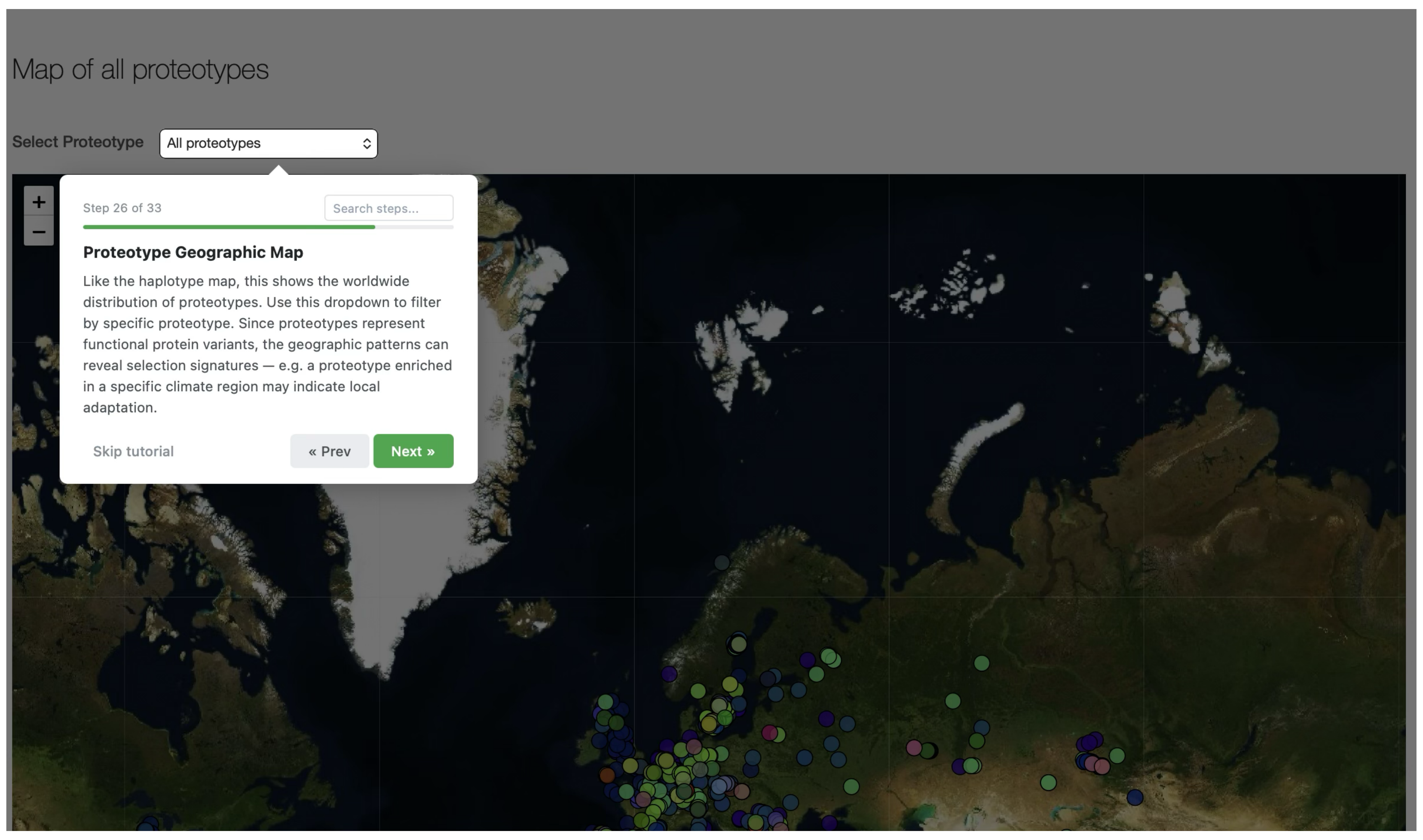

Map of all proteotypes
Select Proteotype
All proteotypes
Step 26 of 33
Search steps...
Proteotype Geographic Map
Like the haplotype map, this shows the worldwide distribution of proteotypes. Use this dropdown to filter by specific proteotype. Since proteotypes represent functional protein variants, the geographic patterns can reveal selection signatures — e.g. a proteotype enriched in a specific climate region may indicate local adaptation.
Skip tutorial
« Prev
Next »


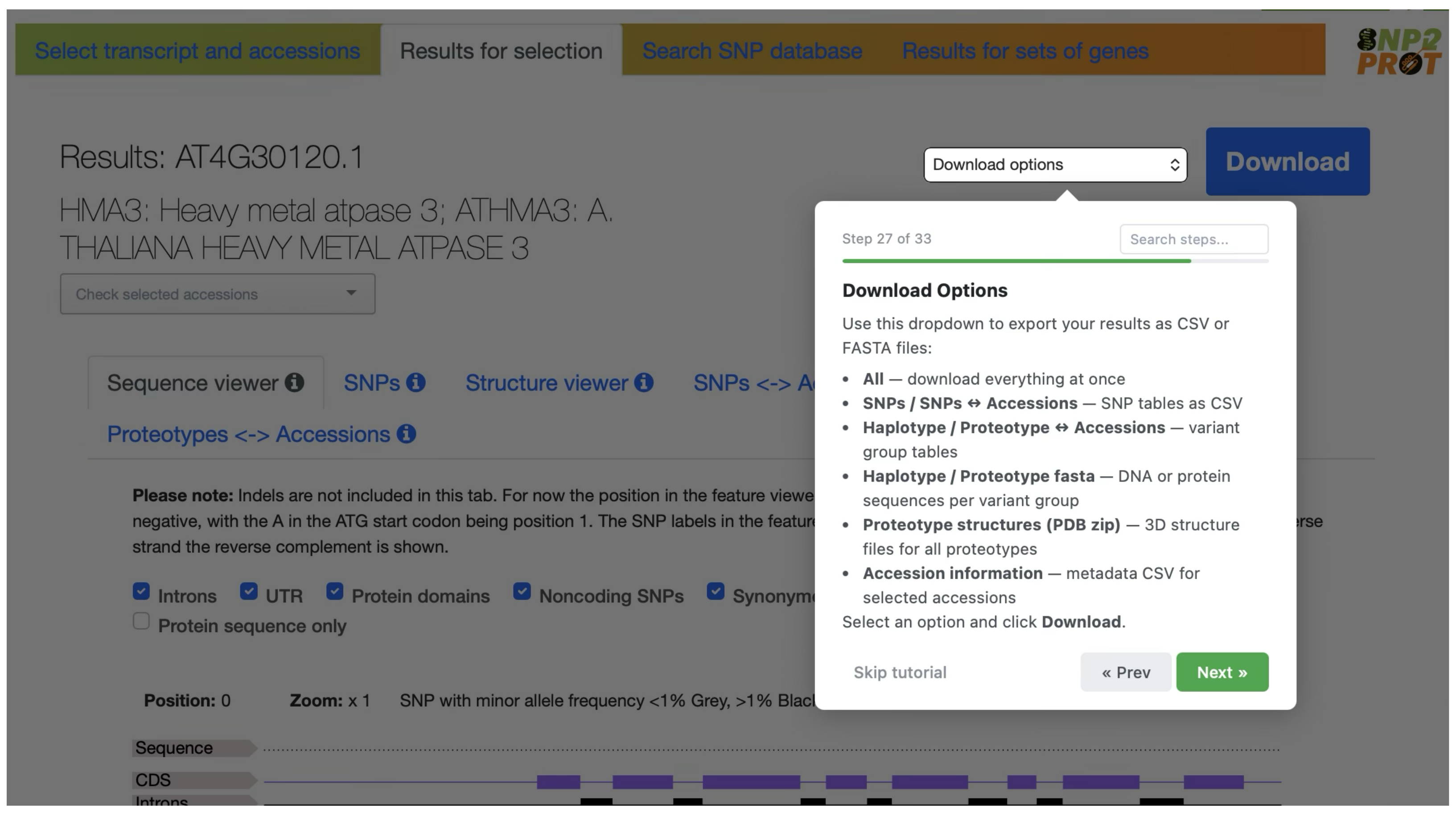

Select transcript and accessions
Results for selection
Search SNP database
Results for sets of genes
Results: AT4G30120.1
HMA3: Heavy metal atpase 3; ATHMA3: A. THALIANA HEAVY METAL ATPASE 3
Download options
Download
Sequence viewer
SNPs
Structure viewer
Proteotypes <-> Accessions
Introns
UTR
Protein domains
Noncoding SNPs
Protein sequence only
Position: 0
Zoom: x 1
Step 27 of 33
Search steps...
Download Options
Use this dropdown to export your results as CSV or FASTA files:
• All — download everything at once
• SNPs / SNPs ↔ Accessions — SNP tables as CSV
• Haplotype / Proteotype ↔ Accessions — variant group tables
• Haplotype / Proteotype fasta — DNA or protein sequences per variant group
• Proteotype structures (PDB zip) — 3D structure files for all proteotypes
• Accession information — metadata CSV for selected accessions
Select an option and click Download.
Skip tutorial
« Prev
Next »


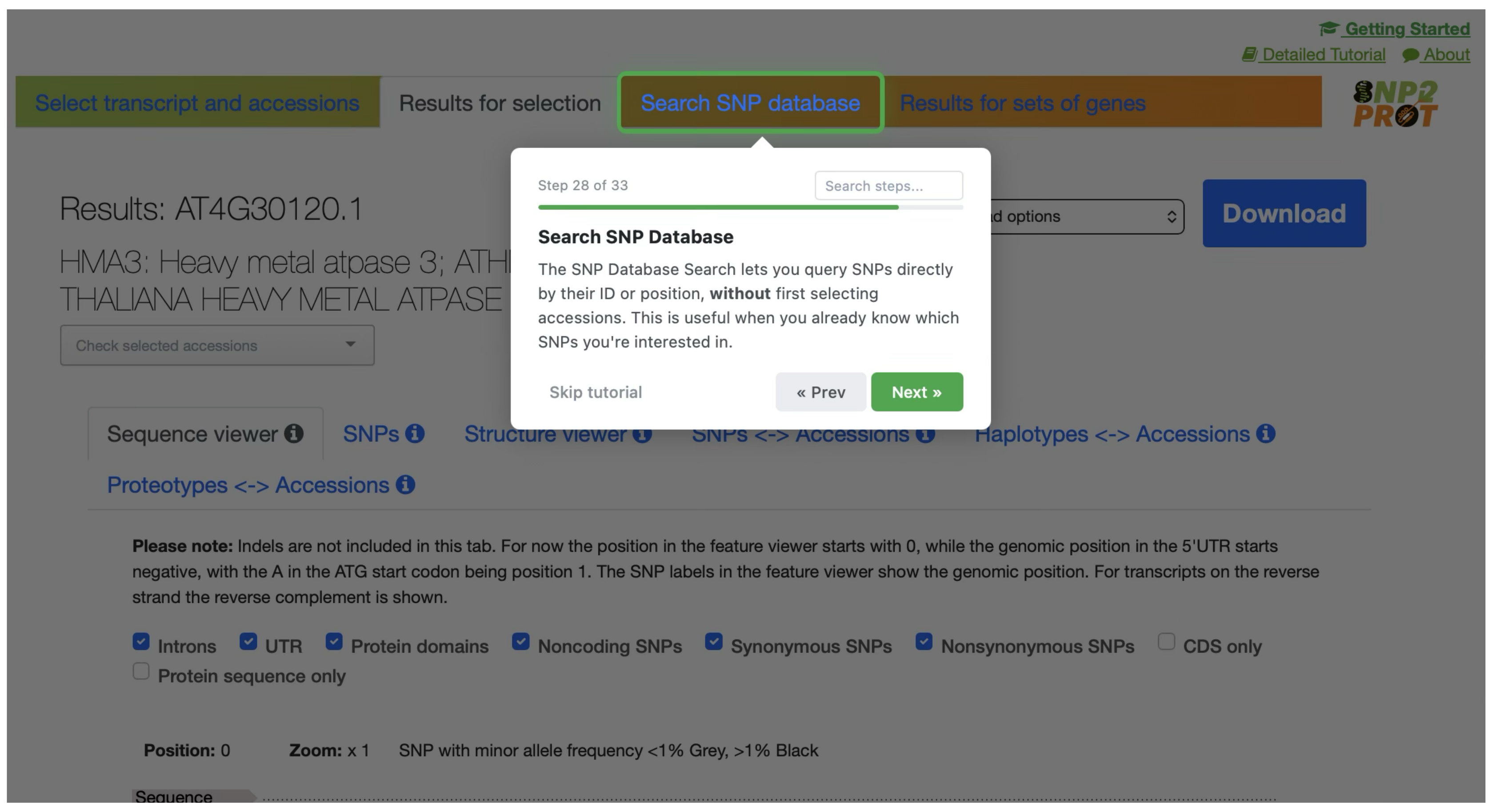

Getting Started
Detailed Tutorial
About
Select transcript and accessions
Results for selection
Search SNP database
Results for sets of genes
Results: AT4G30120.1
Download
Sequence viewer
SNPs
Haplotypes <-> Accessions
Proteotypes <-> Accessions
Please note: Indels are not included in this tab. For now the position in the feature viewer starts with 0, while the genomic position in the 5'UTR starts negative, with the A in the ATG start codon being position 1. The SNP labels in the feature viewer show the genomic position. For transcripts on the reverse strand the reverse complement is shown.
Introns
UTR
Protein domains
Noncoding SNPs
Synonymous SNPs
Nonsynonymous SNPs
CDS only
Protein sequence only
Position: 0
Zoom: x 1
SNP with minor allele frequency <1% Grey, >1% Black
Step 28 of 33
Search steps...
Search SNP Database
The SNP Database Search lets you query SNPs directly by their ID or position, without first selecting accessions. This is useful when you already know which SNPs you're interested in.
Skip tutorial
« Prev
Next »

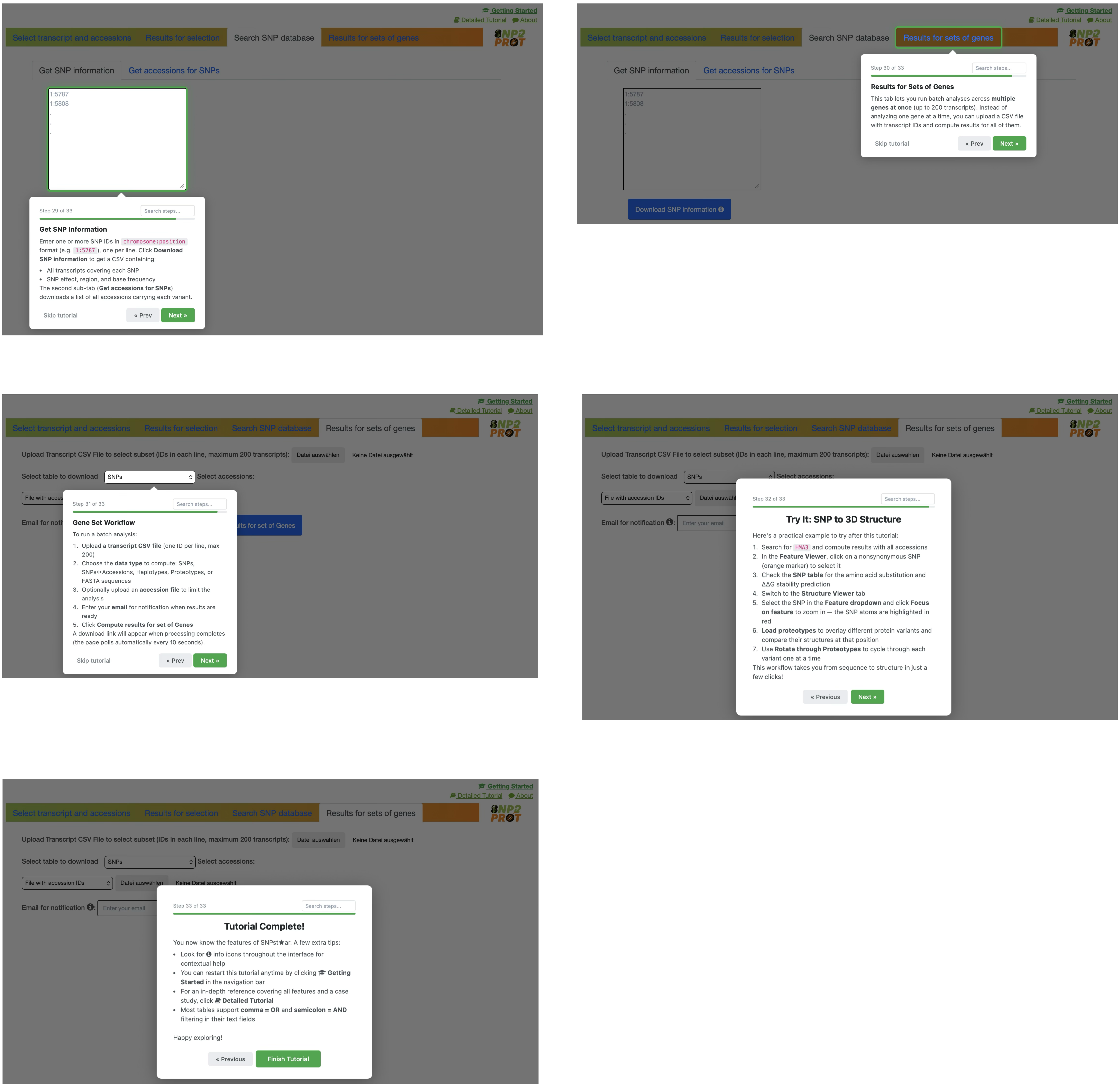


**Figure S1**: Basic step-by-step tutorial introducing SNPstar's features, accessible via the *Getting Started* link on the SNPstar homepage. Tutorial steps are arranged sequentially from left to right and top to bottom. A more comprehensive 61-step walkthrough is available via the *Detailed Tutorial* link.
