## Supplemental Figure 2 for "SNPstar: A Web Server Linking Allelic Variation to Protein Structure and Function in *Arabidopsis thaliana*"

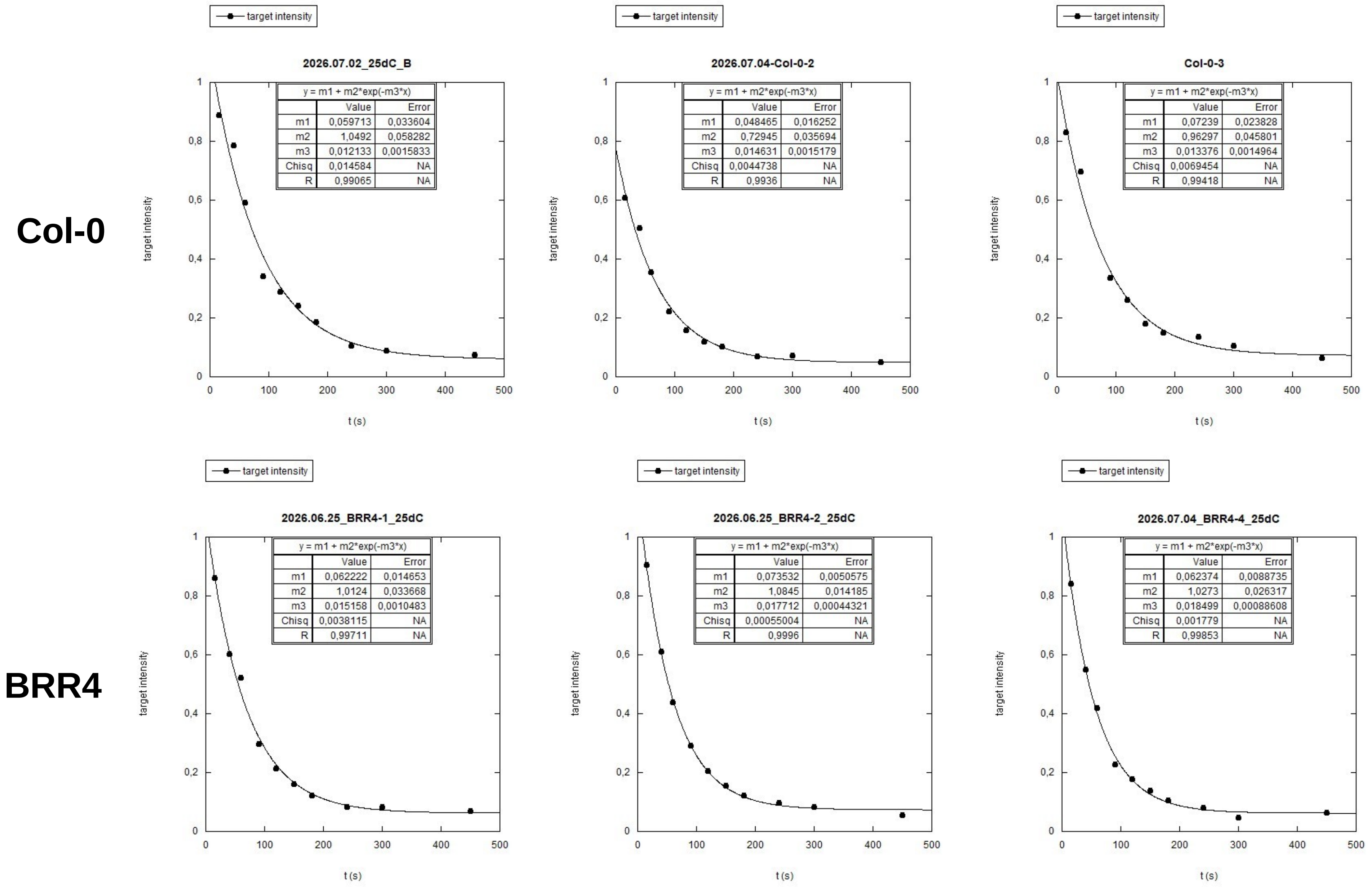


**Figure S2. Single-turnover cleavage kinetics of Col-0 and BRR4 AGO2 in three independent replicates.** *In vitro* slicer assays were performed at 25 °C with FLAG-tagged AGO2/RISC programmed with a GFP-specific siRNA and a fluorescently labelled GFP target RNA. Target-RNA intensity (relative units) was quantified at successive time points and plotted against time (s). Each panel shows one independent experiment for Col-0 (top row) or BRR4 (bottom row). Data were fitted to a single-exponential decay, $y = m1 + m2 \cdot \exp(-m3 \cdot t)$, where *m1* is the non-cleavable plateau, *m2* the amplitude of the cleaved fraction, and *m3* the cleavage rate constant *k* ($s^{-1}$); the fitted parameters, their standard errors, $\chi^2$ and correlation coefficient (*R*) are given in each panel. The per-replicate rate constants were averaged to yield the mean degradation rate constants reported in the main text (Col-0, $k = 0.0134 \pm 0.0013$ $s^{-1}$; BRR4, $k = 0.0171 \pm 0.0017$ $s^{-1}$; mean ± SD, $n = 3$).
